# Structure and assembly of an extremely long bacteriophage tail tube

**DOI:** 10.1101/2022.10.03.510161

**Authors:** Emily Agnello, Joshua Pajak, Xingchen Liu, Brian A Kelch

**Author notes:** Correspondence, address: 364 Plantation St., Worcester MA 01605.

## Abstract

Tail tube assembly is an essential step in the assembly of long-tailed bacteriophages. Limited structural and biophysical information has impeded an understanding of assembly and stability of their long, flexible tail tubes. The hyperthermophilic phage P74-26 is particularly intriguing as it has the longest tail of any known virus (nearly 1 μm) and is the most stable known phage. Here, we present the structure of the P74-26 tail tube and introduce an *in vitro* system for studying the kinetics of tube assembly. Our high resolution cryo-EM structure provides insight into how the P74-26 phage achieves its flexibility and thermostability through assembly of flexible loops into neighboring rings through tight “ball-and-socket”-like interactions. Guided by this structure, and in combination with mutational, light scattering, and molecular dynamics simulations data, we propose a model for the assembly of conserved tube-like structures across phage and other entities possessing Tail Tube-like proteins. Our model proposes that formation of a full ring licenses the adoption of a tube elongation-competent conformation among the flexible loops and their corresponding sockets, which is further stabilized by an adjacent ring. Tail assembly is controlled by the cooperative interaction of dynamic intra- and inter-ring contacts. Given the structural conservation among tail tube proteins and tail-like structures, our model can explain the mechanism of high-fidelity assembly of long, stable tubes.

## INTRODUCTION

Bacteriophages are ubiquitous viruses that selectively and specifically infect their bacterial host. The overwhelming majority of phages are of the order *Caudovirales*, which consist of an icosahedral capsid that contains the double-stranded DNA genome, and a tail. The tail is essential for host recognition and viral attachment, and therefore successful infections, because it serves as the conduit through which the genome travels from the capsid to the host. Tails have morphologically distinct features that further sub-classify them into three families: 1) the short-tailed *Podoviridae*, 2) the long, contractile-tailed *Myoviridae*, and 3) the long, noncontractile-tailed *Siphoviridae*. The tail tubes of long-tailed phage (~85% of all phage) share a common architecture and conserved constituent proteins, suggesting similar principles underlie the assembly and stability of both classes of tails. These tails minimally consist of a Tape Measure Protein (TMP), a complex of tail tip proteins thought to initiate tail assembly, and the Tail Tube Protein (TTP) that polymerizes to form the majority of the tube architecture (1–4). There is clear shared homology and evolutionary origin between TTPs of long-tailed phage, proteins of the bacterial type VI secretion system (T6SS), and bacteriocins (5, 6).

Owing to their long, flexible nature, structural information on tail tubes of siphophages has primarily been limited to pseudo-atomic models using monomeric crystal or NMR structures fit into low-resolution cryo-EM density (7–12). More recently, however, cryo-EM studies have begun to elucidate how TTP is organized in tail tubes (13–15), revealing a conserved fold for TTP, with helically stacked hexameric rings creating a tube whose lumen is occupied by TMP. However, the lack of high-resolution structural information of assembled tubes, combined with a lack of studies revealing assembly kinetics, has limited our understanding of tail tube assembly. To understand the assembly and stability of tail-like structures of phage and other phage-related entities, critical questions regarding conformational changes, biochemical interactions, assembly kinetics, and assembly fidelity remain.

While most siphovirus tails range in length from about 50 to 200 nm (5), a hyperthermophilic phage called P74-26 stands out with the longest tail of any known virus at almost a micron in length (Figure 1A). P74-26 infects the gramnegative bacterium *Thermus thermophilus*, which grows at an optimal temperature of 65 °C (16–17). Owing to the extreme conditions that this phage must endure, it has been characterized as the most stable Caudovirus known (18). Here, we report a 2.7 Å structure of the P74-26 tail tube using cryo-EM. We find that the P74-26 TTP forms rings that are trimeric rather than hexameric, and assembles using an abundance of hydrophobic and electrostatic interactions. Purified P74-26 spontaneously forms flexible tubes that are nearly structurally identical to tails of intact phage. Equipped with the ability to reconstitute tube assembly *in vitro*, we probed protomer and ring interactions through kinetic experiments, mutational analysis, and molecular dynamics simulations to propose a mechanism for tail tube assembly. We find that assembly is governed by formation of ball-and-socket joints, which results in cooperative formation of intra- and inter-ring interactions that overcome autoinhibitory barriers in the monomeric TTP. We propose a model for formation and growth of tail tubes that can explain the high-fidelity mechanism for assembly of long, stable tubes.

**Figure 1.**
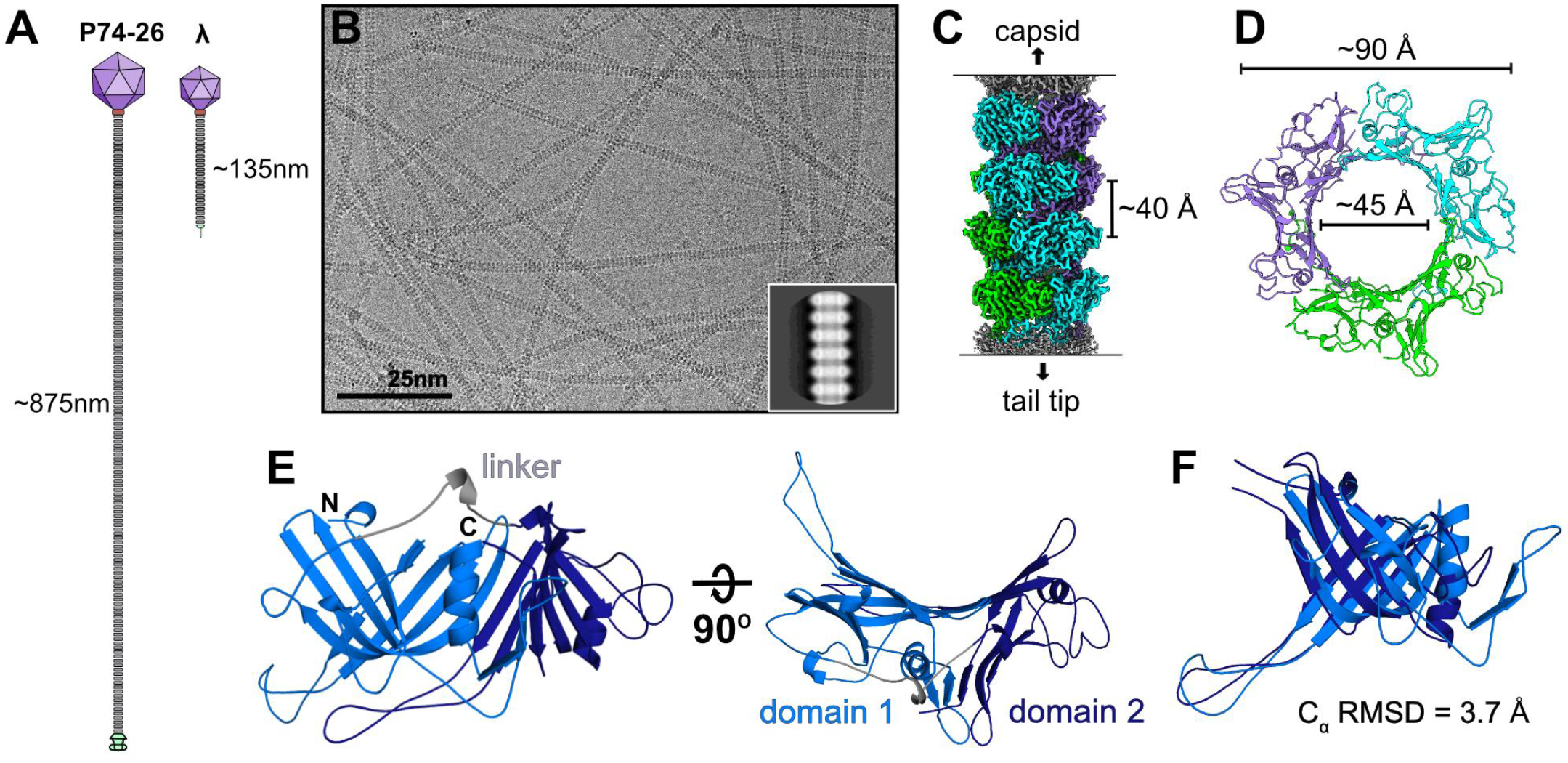
P74-26 tail tube structure. **A** Schematic of siphoviruses P74-26 (left) and λ (right) illustrating the stark difference in tail length. **B** Representative cryo-EM micrograph of P74-26 virions, with an example 2D class (inset). **C** Cryo-EM reconstruction of a segment of tail tube, with four stacked rings highlighted. **D** A single trimeric ring viewed top-down. **E** TTP domain architecture. Each subunit consists of two structurally similar domains. Domain 1 (light blue) is connected to domain 2 (dark blue) by a linker (gray). **F** Overlay of the two domains showing their structural similarity.

## RESULTS

### Structure of the P74-26 virion tail tube

#### Overall Architecture

We determined the structure of the P74-26 tail tube using cryo-EM (PDB ID 8ED0). Micrographs of purified P74-26 virions revealed flexible tails with a stacked ring pattern (Figure 1B). Because of the flexible nature of the tails, traditional single-particle analysis could not be used; instead, we picked sections of tail with ~6 rings per “particle” and used a segmented helical reconstruction approach with the helical symmetry tools in Cryosparc (19) (Supplementary Figure 1). Iterative symmetry searches and helical refinements revealed that the P74-26 tail tube has an overall C3 symmetry, with 3 protomers per ring. C3 symmetry has been seen twice before in the tail tubes of T5 and ΦCbK phage (9, 20, 21). The tube is helical, with a rise of 40.25 Å and a twist of −44.75° (Figure 1C, Supplementary Figure 1. The left-handed helical twist is rare for long-tailed phages, as the majority of tail tubes are right-handed (22, 23). Our final reconstruction has a global resolution of 2.72-Å, according to the gold standard 0.143 Fourier Shell Correlation (FSC) criteria (Supplementary Figure 2A, 2C).

Using this reconstruction, we built an atomic model of the gp93 tail tube protein (TTP) *de novo* in its entirety. We then fit twelve copies of the TTP atomic model into the density of the central four of the six rings in the reconstruction (the two remaining rings at the periphery of the map are much lower resolution). The diameters of the outer and inner surface of the tube are ~90 Å and ~45 Å, respectively (Figure 1D), consistent with other known tail tubes. In some phages, the tail tube has exterior protrusions, such as the immunoglobulin-like (Ig-like) domains seen in T5, λ, YSD1, and Araucaria (9, 10, 13, 24). The P74-26 tail tube, however, has a markedly smooth outer surface with no additional domains or protrusions (Figure 1C). There is density that runs through the center of the tube, which presumably corresponds to the tail’s Tape Measure Protein (TMP), gp95 (Supplementary Figure 2D). The local resolution for this density in the lumen is much lower than the rest of the map, likely due to the segmented method used for reconstruction. We attempted to further resolve the secondary structure in the TMP density, but were unsuccessful and left this density unmodeled. A closer look at the surface of the tail tube lumen reveals surprisingly net neutral electrostatics (see Supporting Text and Supplementary Figure 3)

Each P74-26 TTP subunit consists of two β-sandwich domains: an N-terminal domain (domain 1), and a C-terminal domain (domain 2), connected by a linker. Each domain has the same β-sandwich fold as other tail tube proteins, indicating that it is evolutionarily related to other TTPs despite the difference in overall symmetry. Much like single-domain TTPs, each β-sandwich domain contains a long hairpin that emanates from the ‘bottom’ of the TTP subunit; we term these loops “Loop1” (from domain 1) and “Loop2” (from domain 2) (Figure 2C). The two domains are structurally similar to each other (C_α_ RMSD of ~3.7 Å over 113 out of 174 residues; Figure 1F), despite having very little sequence similarity (16% identity). Therefore, the two-domain architecture of P74-26 TTP likely arose through an ancient gene duplication and fusion event. Thus, despite the C3 symmetry, the overall ring can be considered pseudohexameric, explaining the similarity to hexameric tail tubes of most long-tailed phages. Furthermore, the N-terminal domain of one subunit is related to the C-terminal domain of a subunit in an adjacent ring in a right-handed fashion with a twist of ~19°, consistent with the helical twist of right-handed hexameric tail tubes. Similar pseudo-symmetry is also seen in siphophage T5, where each subunit similarly consists of two β-sandwich domains (9). As in T5, each P74-26 TTP domain is structurally similar to the TTP of other phage and R-pyocins, illustrating a shared evolutionary lineage (5, 6, 9).

**Figure 2.**
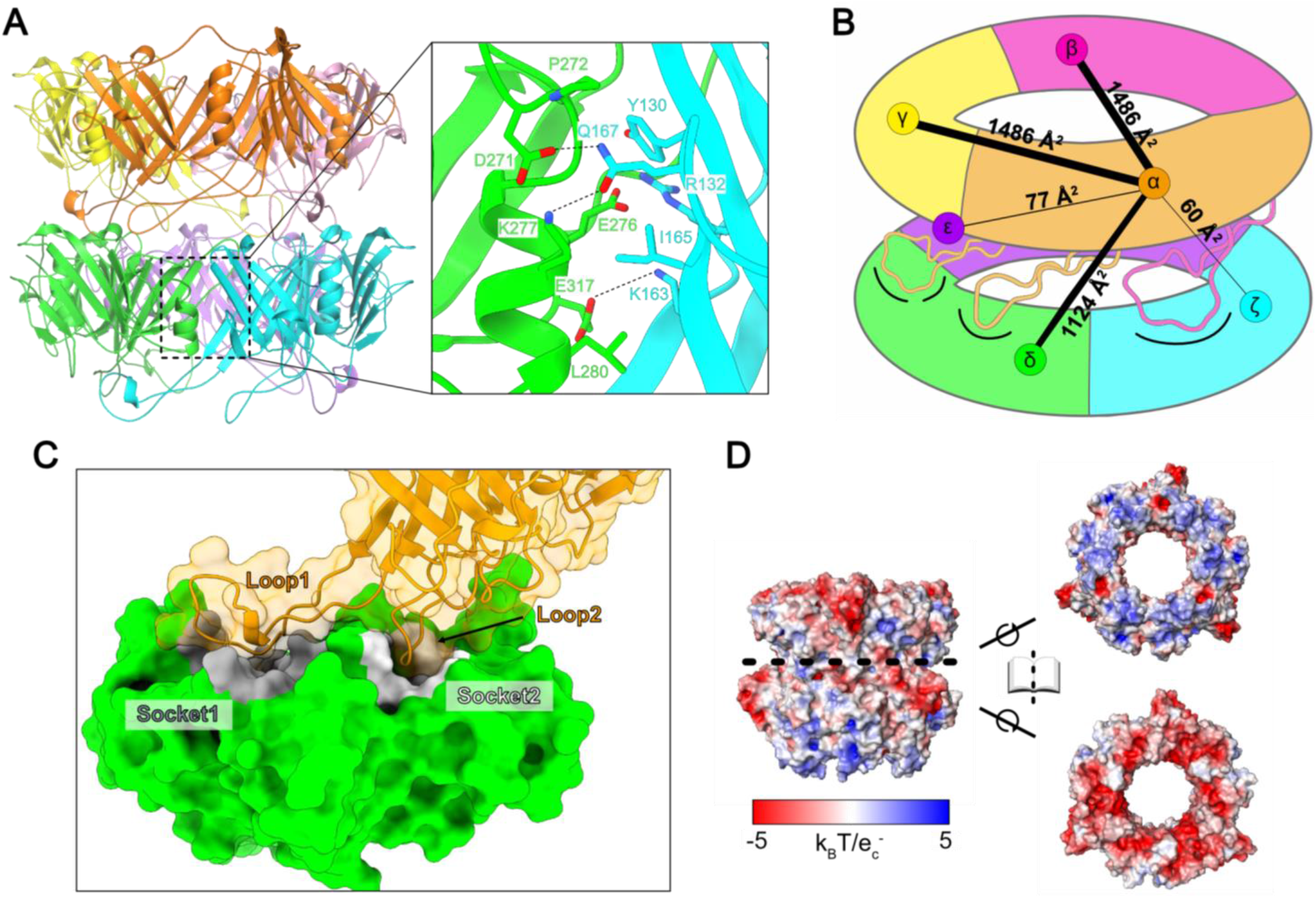
Intra- and inter-ring interactions. **A** The primary intra-ring interface between two subunits (green and cyan) within a ring. **B** Schematic quantifying interactions from a single subunit (α) to illustrate the extensive, cooperative network. Numbers and line widths correspond to quantification of the interactions between α and neighboring subunits as calculated by the PDBePISA server (Supplementary Tables 3 and 4). **C** Surface representation of two subunits reveals a ball and socket geometry between rings. A single subunit (orange) has two loops (Loop1 and Loop2) that fit into sockets (Socket1 - gray, and Socket2 - white) of a subunit in the ring below it (green). **D** Surface electrostatics of ring interfaces demonstrate an important role for electrostatics in inter-ring interactions.

#### Inter- and Intra-ring interactions

The tail tube is held together by two types of interfaces: within a ring (intra-ring) and between rings (inter-ring). Within each ring, subunits are arranged head-to-tail into a trimeric ring, reminiscent of DNA polymerase sliding clamps (ring-shaped proteins that also evolved through domain duplications) (25–27). Each subunit contacts six subunits inter-ring (three ‘below’ and three ‘above’.) The intra-ring interaction area is slightly larger than the inter-ring (3 x 1486 Å^2^ = 4458 Å^2^ versus 3 x 1261 Å^2^ = 3783 Å^2^; Figure 2B, Supplementary Table 4).

The intra-ring interactions come from two interfaces: a direct interaction between the two β-sandwich domains of adjacent subunits, and another interface between Loop1 of one subunit and the underside of the adjacent subunit’s domain 2 (Figure 2A). Both interfaces are particularly rich in hydrophobic interactions. For example, Met1, Tyr130, and Phe123 in domain 1 bind to Pro272, Ala253, Ala 273, Val322, and Ile328 of domain 2, while Met55 in Loop1 inserts into a hydrophobic pocket formed by Leu108, Val244, Ile308, and Ala336 of domain 2. We further note that the N-terminal methionine directly contributes to intra-ring interactions, providing a structural explanation for previous work showing that the flexible N-terminus controls tail assembly in other phages (8, 10).

Nearly all inter-ring interactions are mediated by Loop1 and Loop2. Loop1 makes slightly more extensive inter-ring contacts than Loop2 (811.9 Å^2^ vs 521.6 Å^2^) (Supplementary Table 4). Thus, Loop1 plays a critical role in stabilizing interactions both within rings and between rings. Each TTP subunit contacts both other subunits within the same ring as well as all subunits in both adjacent rings, creating a cooperative unit (Figure 2B). The inter-ring interactions are mediated by a ball-and-socket-like geometry wherein the tips of Loop1 and Loop2 each fit into a socket of a subunit in the adjacent ring; Socket1 and Socket2, respectively (Figure 2C). Furthermore, this interface is supported by extensive electrostatic interactions, with the surface of the loop-side harboring a net positive charge while the socket-side is negative (Figure 2D). While the overall loop and socket geometry appears to be conserved across tail tubes, the P74-26 tail tube uses exaggerated ionic interactions for enhanced thermostability (see Supporting Text and Supplementary table 3).

### Tail tube protein polymerizes *in vitro*

To investigate P74-26 TTP polymerization *in vitro*, we recombinantly expressed and purified the P74-26 TTP in *E. coli* (Supplementary Figure 4A). When we examined purified, soluble TTP by negative stain EM, we observed long structures resembling tail tubes (Supplementary Figure 4B). These tubes form a range of lengths but are on average much longer than virion tails, presumably because the Tape Measure Protein that regulates tube length during virion formation is absent (28). In addition, the *in vitro*-assembled tubes exhibit greater flexibility than virion tails (Supplementary Figure 4D). Thus, we posit that the Tape Measure Protein contributes to the stiffness of virion tails.

To evaluate the similarity between the *in vitro-* assembled tubes and tails from intact virions, we determined the structure of the reconstituted tubes using cryo-EM (PDB ID 8EDX) to a global resolution of 2.8-Å (Supplementary Figures 2B, 2E and 4C), using a similar helical reconstruction protocol as we implemented for virion tails. We found that *in vitro* assembled tubes are structurally identical to the virion tail structure (Cα RMSD ~ 0.11 Å across all 348 residues), except there is no density for TMP running through the center of the tube, as expected (Supplementary Figure 2E). Therefore, the *in vitro* assembly of TTP into tubes establishes this system as a useful tool for revealing the mechanism of tail tube polymerization.

To understand how TTP polymerization occurs, we asked what the dominant oligomerization state of TTP in solution is. We filtered purified TTP to remove any spontaneously polymerized tubes and analyzed the flow-through using Size Exclusion Chromatography with MultiAngle Light Scattering (SEC-MALS). We observe that the majority of soluble TTP is monomeric in solution (~76% mass fraction; Figure 3A). This observation is not unexpected, as TTP from other phages are primarily monomeric in solution as well (9, 13, 29). We also observe a minor peak whose molecular weight is consistent with a hexamer of TTP (mass fraction ~6%).

**Figure 3.**
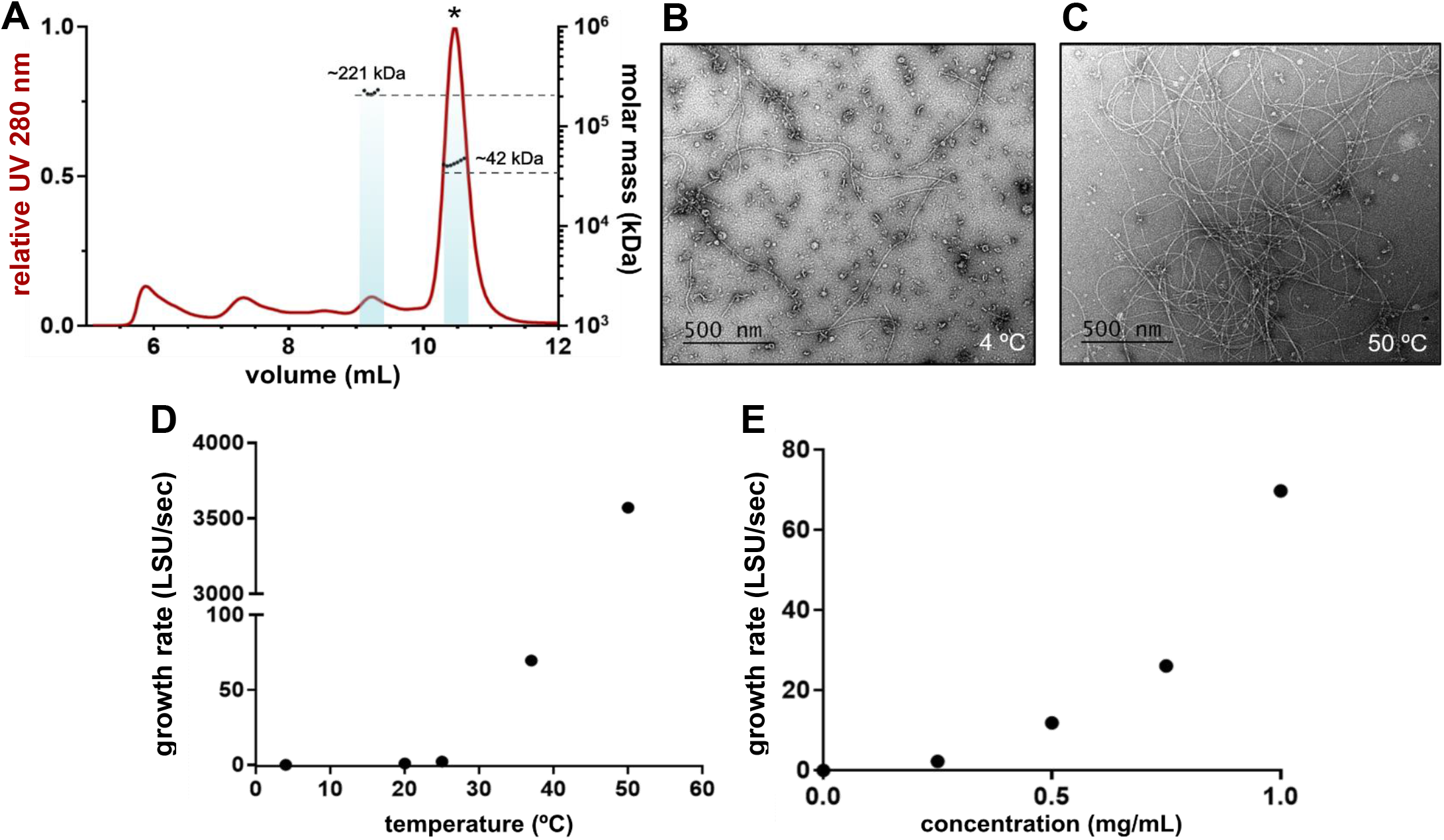
Properties of *in vitro* TTP assembly. **A** SEC-MALS of filtered TTP, displaying the UV trace (red) and calculated molecular weights (black dots) with an average calculated molecular weight of 42.3 ± 0.347 kDa and 221 ± 8.18 kDa 3.7%. Dashed lines display the true molecular weights for a monomer (~37.7 kDa) and a hexamer (~226 kDa). **B** and **C** Negative stain images of *in vitro-assembled* tubes incubated for 3 hours at 4 °C **(B)** or 50 °C **(C)** indicate tube formation is enhanced by heat. **D** Calculations of growth rates in Light Scattering Units per second, detailed in Supplementary Figures 5 and 6. Tube formation is non-linearly accelerated by higher temperature. **E** Tube growth rates increase non-linearly by higher TTP concentration.

Because of the thermophilic nature of P74-26, we explored the temperature-dependence of our *in vitro-* assembled tube formation. We applied purified TTP to negative stain grids after incubation at 4 °*C* and 50 °*C* for 3 hours. There was an increase in the number of tubes per image as the temperature increased, suggesting tube assembly was promoted by higher temperatures (Figure 3B, 3C).

As sample application in negative stain EM can be variable, we sought to establish a more quantifiable method for measuring tube assembly under a range of conditions. To do so, we took advantage of the mass increase that accompanies tube growth, increasing light scattering as assembly progresses. Using a light scattering assay, we monitored tube assembly from purified TTP at 4 °C, 20°C, 25°C 37°C, and 50°C. Consistent with the negative stain results, tube formation increased with temperature (Supplementary Figure 5A). Kinetically, the assembly of tail tubes over time at 4°C and 20°C progressed linearly, while the 37°C displayed a sigmoidal shape and the 50 °C condition exhibited a fast initial burst of assembly followed by a plateau (Supplementary Figure 5A). To quantitatively compare rates of tube formation across this temperature range, we calculated the rates during the fast-phase of tube growth for each condition (Figure 3D). While assembly of TTP tubes occurred slowly over 3 hours at 20°C, assembly occurred much faster at 37°C and even faster still at 50°C, with rates ~68x and 3,516x the 20°C rate, respectively (Supplementary Figure 5C). The relationship between temperature and growth rate is non-linear and demonstrates a steep temperature-dependence, suggesting there is an entropic barrier to assembly.

Using this same assay, we then measured the concentration-dependence of tube formation at 37 *C*. The curves of light scattering versus time were sigmoidal in shape (Supplementary Figure 6A), so we again focused on the fast rate of the curves confirming that increased concentrations result in a faster rate of assembly (Supplementary Figure 6D). We then plotted the rates of assembly against TTP concentration and found that the concentration dependence is non-linear (Figure 3E). This indicates a complex relationship with TTP concentration and suggests that initial seeding and tube elongation may proceed with different kinetics.

### Mutational analysis of tube formation

Using our cryo-EM structures, we hypothesized that the intra- and inter-ring interactions control tube assembly. We tested whether these interactions are important for tube growth using site-directed mutagenesis and *in vitro* tube growth assays. To target intra-ring interactions, we generated two separate variants: the Y130A mutation (Y130 sits in a hydrophobic pocket at the interface between two subunits) and the Q167A mutation (Q167 hydrogen bonds to the adjacent subunit) (Figure 4). To target inter-ring interactions, we generated four separate variants (Figure 4). The L229A mutation tests the role of L229, which sits at the tip of Loop2 and interacts with the corresponding socket on the subunit of the next ring through hydrophobic interactions. Because there aren’t any residues in domain 1 that make as extensive contacts as L229, we made quadruple alanine mutations at the tip of Loop1 (48-RAIRR-52 to 48-AAAAA-52; termed the Loop1-Ala variant). We also sought to disrupt the sockets, by mutating the loop that comprises the wall between the two sockets from 147-RVNDM-151 to 147-AAAAA-151 (termed the socket-Ala variant). Finally, to test the role of Loop1 and if assembly could occur with only one loop per subunit, we deleted the entire Loop1 (Q43-T65; the Δloop1 variant).

**Figure 4.**
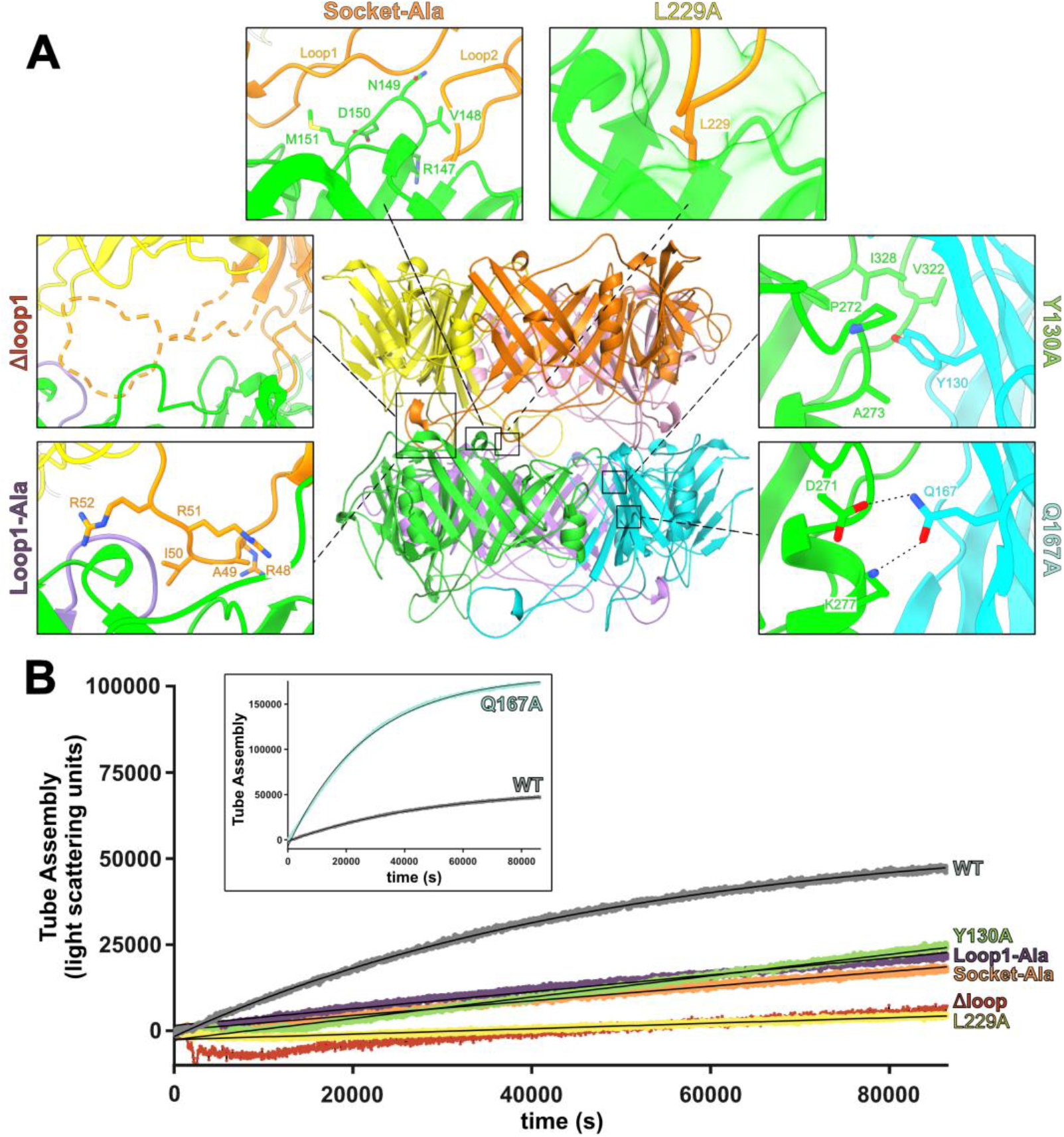
*In vitro* mutational analysis of tube assembly. **A** Location of the chosen TTP mutations in the context of the tube (center), and zoomed-in views of residues selected for mutation. **B** Light scattering curves of TTP variant tube formation at 1 mg/mL over time, with higher light scattering intensity values (counts per second) indicating a more highly assembled state. Variant Q167A suggests a negative regulatory role for residue Q167 in tube formation (inset). Negative stain EM confirming assembly and a summary of rates can be found in Supplementary Figure 8.

TTP variants were purified and confirmed to be primarily monomeric in solution after filtering (Supplementary Figure 7). After filtering, we monitored tube formation by light scattering. We note that these experiments were done at 20 *°C* rather than at 37 *°C* to minimize any complications from potential thermostability defects. The Δloop1 variant had drastic effects, with light scattering signal just barely above zero. Negative stain EM confirmed almost no tubes, but with the very occasional presence of short tubes, suggesting the variant is folded and capable of assembly, but that assembly is severely disfavored (Supplementary Figure 8G). This suggests that Loop1 is required for tube assembly. The Loop1-Ala variant, however, had tube formation similar to WT, based on both light scattering and negative stain, suggesting that the necessary interactions within this loop are not specific to these inter-ring residues (Figure 4B, Supplementary Figure 8D). In contrast, Loop2 relies heavily on a single interaction: the L229A residue at the tip of the loop which sits in a hydrophobic pocket in Socket 2. The L229A variant severely limited the rate and extent of tube formation, with a tube formation rate similar to that of the Δloop1 variant (~10% of the WT), but with tubes still sparsely distributed in negative stain EM (Figure 4B, Supplementary Figure 8F). Similarly, the socket-Ala variant also limited tube assembly with far less tube formation than WT (~18% of the WT rate), but not completely abrogated (Supplementary Figure 8E). The intra-ring mutations had varying results with the Y130 residue appearing to be important to tube formation while the Q167A variant had increased levels of tube formation, suggesting the Q167 residue may play an inhibitory or regulatory role in assembly (Figure 4, Supplementary Figure 8B). Variant assembly rates are summarized in Supplementary Figure 8H. This mutational analysis reveals that while intra-ring interactions are important, tube growth is critically dependent on inter-ring interactions.

### Dynamics of TTP oligomers

Because tail assembly is a dynamic process, we wanted to investigate the behavior of TTP prior to tube assembly. To yield insight into the preferred conformations of monomeric TTP, we performed four independent ~2 μs MD simulations using a single subunit from our tail structure as the starting model. During the simulations, the β-sandwich domains remain relatively stable. However, Loop1 and the N-terminal residues exhibit significant flexibility and conformational changes (Figure 5A and 5D). Loop1 is highly flexible, permitting it to fold back and interact with Loop2 and the β-sandwich domain, where it is stabilized (Figure 5D and Supplementary Figure 9). Further, this flexibility allows it to form secondary structural motifs (Supplementary Movie 1). Thus, the conformational ensemble of Loop1 in soluble monomers is largely incompatible with the intra- or inter-ring binding modes seen in the assembled tube. Moreover, Socket1 can become blocked by the N-terminal methionine, which flips ~180° to interact with Phe33. This steric block is further stabilized by Arg2 interacting with an acidic patch consisting of Asp184, Glu185, and Glu186. Thus, we predict that the monomer state inhibits assembly due to the flexibility of Loop1 and the closing of Socket1.

**Figure 5.**
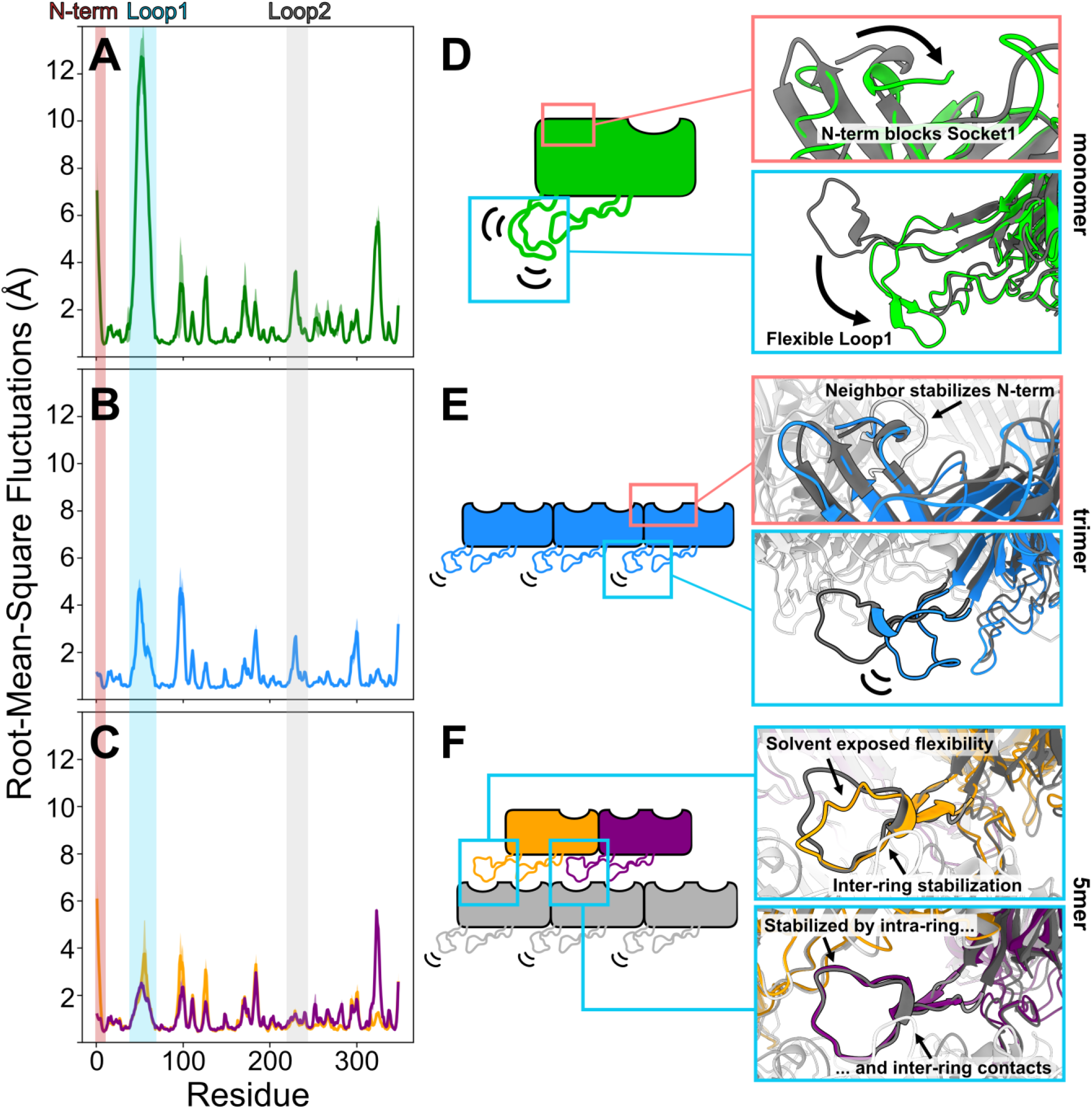
Progressive stabilization predicted by molecular dynamic simulations. **A-C** RMSF plots of monomer **(A),** trimer **(B),** or 5mer **(C)** simulations. Mean values across all replicates are lines colored according to the schematic figures shown in **D-F**, and the standard error in the mean is colored in a lighter shade. Shaded regions correspond to the N-term (red), Loop1 (cyan), and Loop2 (gray) to highlight the stabilization of these regions at different stages of assembly. **D-F** Cartoon schematics of predicted dynamics with insets visualizing simulated structures (color) superimposed onto the cryo-EM reconstruction (grayscale). The simulated structures are representative centroids of the largest cluster of a *k*=10 means clustering of an individual simulation. These structures are provided as PDB files in the supplemental information.

To see how these auto-inhibitory interactions are altered upon ring formation, we simulated a trimer ring. These simulations reveal that intra-ring interactions poise the trimer for tube elongation by alleviating some of the inhibitory conformations that prevent inter-ring assembly in the monomeric state. In the trimeric state, the Met1 sidechain forms part of the hydrophobic interface between subunits, locking the N-terminus into the conformation seen in tubes, with no significant blocking of Socket1 (Figure 5B, 5E). Therefore, the intra-ring interactions “lock” Socket1 into an open conformation that is competent for binding of future inter-ring interactions. Similarly, Loop1 is also partially stabilized by intra-ring contacts (Fig. 5B). However, the loop remains somewhat flexible. This flexibility ultimately results in a loss of native contacts during the simulations between Loop1 and the neighboring subunit (Fig. 5E, Supplementary Figure 10, and Supplementary Movie 2). The linchpin for Loop1 seems to be hydrophobic interactions between the top of the loop and a hydrophobic pocket of the neighboring intra-ring subunit; when this interaction is lost, Loop1 becomes much more flexible (Supplementary Movie 2). Therefore, the formation of a trimeric ring positions Socket1 into a competent conformation, but Loop1 remains too flexible to support inter-ring contacts.

To investigate the role of both intra- and inter-ring contacts in potential assembly intermediates, we simulated a pentameric arrangement: a trimeric ring with two subunits of the ring above. This subunit arrangement allows us to examine how Loop1 responds to different environments, as we have a single Loop1 that is making only inter-ring contacts, one Loop1 that is making both intra- and inter-ring contacts, and three Loop1s that only make intra-ring contacts. In our pentamer simulations, Loop1 is considerably more rigid when it is stabilized by *both* intra-ring and inter-ring interactions (Figure 5C). Interestingly, Loop1 appears to be more stabilized by inter-ring interactions alone than intra-ring interactions alone (Figure 5B and 5C). An important distinction is that the flexibility of the loop making only inter-ring interactions is primarily in the solvent exposed region (Figure 5F), and this loop maintains native contacts better than loops stabilized by only intra-ring contacts (Supplementary Figure 10). Together, these simulations indicate that the vertical stacking of subunits is largely responsible for the rigidification of Loop1. Furthermore, our pentamer simulations also yield insight into how an incomplete ring could accept the final subunit (Supplementary Figure 11 and Supplementary Movie 3). Altogether, our simulations indicate oligomers smaller than a hexamer (two stacked rings) are not stable due to flexible conformations of Loop1 and Socket1. We note experimental support for this prediction, as our SEC-MALS data shows a stable hexamer in solution (Figure 3A). Thus, our simulations of a TTP pentamer illustrate how the sockets and loops are linked to cooperative formation of intra- and inter-ring interactions.

Finally, we sought to ascertain if Loop1 and Socket1 conformationally control assembly of TTPs from other phages. We chose to examine the TTPs of two siphophages, SPP1 and YSD1, because structures are available (12, 13). Further, this a good test for the generalization of our findings, as the SPP1 and YSD1 tail tubes consist of true hexameric rings composed of single-domain subunits rather than the trimeric rings of two-domain subunits of P74-26. First, we performed MD simulations of TTP^SPP1^ and TTP^YSD1^ in their monomeric states. Like our TTP^P74-26^ monomer simulations, these simulations predict highly flexible N-terminal and Loop residues (Supplementary Figures 12 and 13, and Supplementary Movies 4 and 5). In these simulations, the area that contributes to formation of the socket can be blocked by the N-terminal residues and, in the case of YSD1, additionally blocked by the Ig-like domain (Supplementary Movies 4 and 5). Further, our simulations also predict that the loop (corresponding to Loop1 of P74-26) of TTP^SPP1^ and TTP^YSD1^ folds back into a conformation incompatible with assembly (Supplementary Figures 12 and 13).

We next performed MD simulations of TTP^SPP1^ and TTP^YSD1^ in an arrangement of two rings minus a single subunit, similar to our 5mer simulations of TTP^P74-26^. Like our TTP^P74-26^ pentameric simulation, these simulations suggest that the N-terminal residues (and also the YSD1 Ig-like domain) are rigidified upon assembly, promoting open socket interactions for future inter-ring contacts. Further, the Loop is stabilized by inter-ring and intra-ring contacts alike. In the case of YSD1, the trend of Loop stabilization follows that of P74-26, where loops stabilized by both inter- and intra-ring contacts are most stable, and only inter-ring contacts provide slightly more stabilization to the loop than only intra-ring contacts (Supplementary Figure 14 and Supplementary Movie 6). On the other hand, our TTP^SPP1^ 11mer assembly is still highly flexible and preserves little-to-no native contacts between the loop and neighboring subunits (Supplementary Figure 15 and Supplementary Movie 7), suggesting that there may be additional considerations for this assembly. Our simulations of TTP^SPP1^ and TTP^YSD1^ provide supportive evidence that the mechanisms underlying the predicted P74-26 tail tube assembly model are largely conserved across long-tailed phage.

## DISCUSSION

### Mechanisms for cooperative tube assembly

To form tail tube assemblies, individual subunits make extensive interactions within rings and between rings. Within a ring, subunits interact through extended beta sheets and contacts made at Loop1. Between rings, subunits assemble through complementary loop/socket geometry, where the loops of one ring fit snugly into the sockets of an adjacent ring. Notably, this indicates that Loop1 is important for both intra-ring and inter-ring assembly, consistent with the deletion of this loop completely abrogating assembly *in vitro* (Figure 4). This highly interwoven network of interactions likely enhances the stability of the entire assembly in a cooperative manner.

How this kind of interlocking network arises from monomer self-assembly is not obvious. As with all phage tails, defects in tail tube assembly would compromise the stability of the tail and ultimately compromise productive phage infection. Because the P74-26 tail tube is exceptionally long, the need to avoid off-pathway assembly is exaggerated, as low-probability defects have more opportunities to arise. Thus, mechanisms that prevent off-pathway assemblies and guide monomers to correct configurations are required, and these mechanisms are likely conserved across many siphophage tail tubes.

Such mechanisms likely regulate each step of self-assembly, including the assembly of monomeric building blocks into trimeric rings, as well as the assembly of rings to higher-order structures. We will therefore discuss assembly mechanisms at the level of a monomer, then in the context of intra-ring interactions, and finally at the level of inter-ring interactions.

### A proposed mechanism for tail tube polymerization

Our high-resolution structure of the P74-26 tail tube reveals specific global and residue-level interactions important for tube assembly. The most basic building block of the tail tube is a monomer, which our MALS data shows is the predominant species in solution (Figure 3A). Because there is a high population of monomers in solution, we anticipated that the monomer would have mechanisms regulating both intra-ring and inter-ring contact formation, which is further supported by the faster tail tube formation we observed at higher temperatures. These data indicate that there is an entropic barrier for monomers to associate into higher-order structures, and that thermal energy helps overcome this barrier. We envision the following non-mutually exclusive mechanisms for regulating this process.

First, our simulations predict that, in the monomeric state, Loop1 can adopt a wide range of meta-stable conformations. Many of these conformations involve Loop1 making contacts with Loop2, precluding both from interacting with other monomers. Because Loop1 is important for both intra- and inter-ring assembly, this mechanism alone can explain an entropic barrier to monomer association; higher temperature can peel Loop1 away from the globular domain and allow it to sample extended states that are competent for binding another subunit’s socket.

Second, we find that intra-ring interactions control the conformation of Socket1, which thereby controls inter-ring interactions. In our simulations of a TTP monomer, we find that Socket1 is sterically occluded with the N-terminal five residues. However, our simulations of a TTP trimer ring show that Socket1 is unblocked because the N-terminus folds back toward the intra-ring interface. This reorganization of the N-terminus is driven by hydrophobic interactions between the N-terminal methionine and hydrophobic residues that participate in the subunit-subunit interface (Phe123 and Tyr130 in *cis*, and Ala253 and Pro270 in *trans*). Because Met1 plays a critical role in forming the intra-ring interface, intra-ring assembly forces the N-terminus to vacate Socket1, which permits Loop1 from another monomer to establish inter-ring contacts. This supports a mechanism by which trimeric rings first form. thereby locking the sockets of all three monomers in a state that promotes inter-ring assembly (Figure 6C). Therefore, the formation of a single ring allows stepwise assembly of the adjacent ring (Figure 6D).

**Figure 6.**
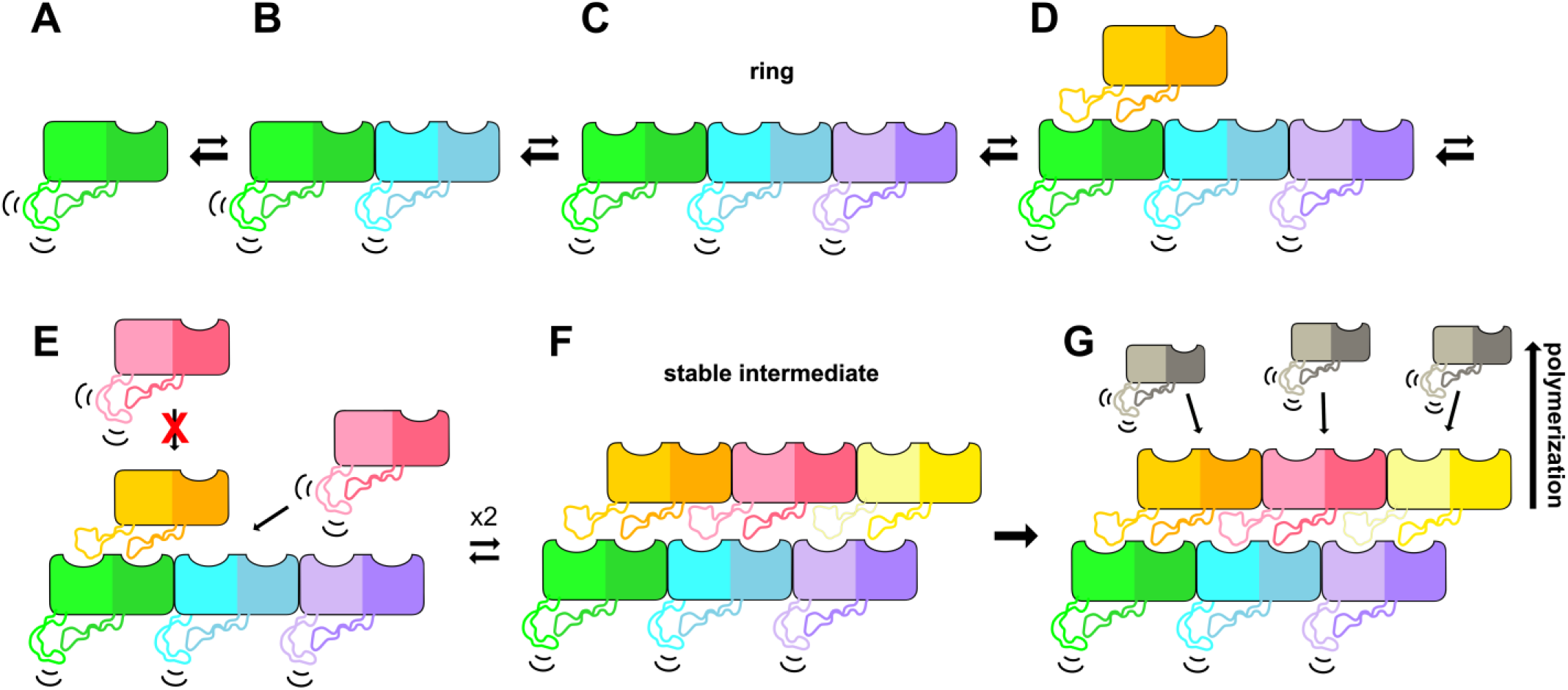
Proposed model for tail tube assembly. **A** Monomeric TTP in solution has a closed Socket1 and flexible Loop1. **B** Addition of a subunit transiently stabilizes Socket1. **C** Trimeric ring formation leads to opening and stabilization of all Socket1 s within the ring, but with flexible Loop1s. **D** Open sockets on the trimer allow for subunit addition. **E** Due to the ball and socket geometry, further subunit addition is likely to occur laterally, completing a ring through the addition of two additional subunits to form **(F). F** A two-ring complex is predicted to be the first stable intermediate, creating a platform for polymerization. **G** Polymerization of the tail tube proceeds spontaneously.

In our *in vitro*-assembly system, we do not observe trimers by SEC-MALS, indicating that the trimer is likely not stable. As discussed, our simulations predict that Loop1 is highly flexible even in a trimer, but this loop is only stabilized by the binding of another ring below it. Therefore, the interplay of inter- and intra-ring interactions would mutually stabilize both rings. Thus, a two-ring complex achieves a highly stable scaffold for polymerization to occur from. In fact, we observe a low population of hexamers by SEC-MALS (Figure 3A). Therefore, we predict that two TTP rings is the primary stable intermediate for subsequent assembly in our *in vitro* assembly reactions (Figure 6F). We expect that *in vivo*, the stabilization of a single ring may be achieved by other factors, such as the Tail Tip Complex (TTC) or Tail Assembly Chaperones. Tail Assembly Chaperones have been shown in phage Λ to be essential for phage tail production and to bind both TTP and TMP (30, 31). Thus, in addition to potentially playing a role in TTP stabilization, they are also a likely candidate for the negative regulation that would need to occur *in vivo* to ensure assembly is occurring in concert with other essential factors like the TMP on a functional timescale.

Once a stable intermediate has formed, tube polymerization is likely to occur by addition of monomers one by one, completing one ring of three subunits before starting to form the next ring (Figure 6G). If the rate-limiting step is formation of a trimeric ring as suggested by our data, it is unlikely that rings would form in solution and then be added ring-by-ring. Rather, monomers are added and thereby stabilized by inter-ring interactions while the intra-ring interactions form as two more subunits fill in the ring. In the context of assembly *in vivo*, this mechanism makes sense for multiple reasons. For one, the tail tube must form around the TMP which creates topological constraints that would make ring addition difficult, akin to threading beads onto a string. Because a monomer does not have its Socket1 stabilized, it is more likely to bind using its loop-face. Thus, our model predicts that tail tubes assemble using a unidirectional polymerization pathway: monomers can only bind to the face of the ring harboring the sockets because the ring’s loops are still flexible. Unidirectional growth could be advantageous in the case of a single polymerization initiation event (at the Tail Tip Complex, for example), from which polymerization extends. Finally, the loop-socket geometry ensures high-fidelity polymerization; it would discourage addition of a monomer to an unfinished ring, preventing aberrant subunit addition and ensuring minimal defects in tube assembly.

The structural conservation between tail tube proteins suggests conserved mechanisms for tube assembly. Our model hinges on the importance of the N-terminus and Loop1 as both autoinhibitory elements in the monomeric state and critical contact points for intra- and inter-ring interactions. Despite the differences in symmetry between P74-26 and other tail tubes, these regions are known to be essential in a number of hexameric systems. In phage λ, a reorganization of the N-terminus and loop occurs between the monomeric and assembled states: the N-terminal region becomes trapped by interactions with the next subunit and the loop is captured along the inter-ring interface (10). These regions are proposed to be critical for tube polymerization in diverse entities including phages T5 and T4, as well as tail-like complexes such as T6SS, pyocins, and extracellular contractile injection systems (5, 8–10, 12, 32). These observations not only validate our proposed polymerization model but also suggest that these mechanisms are used across a diverse range of phage tail and tail-like assembly pathways.

## MATERIALS AND METHODS

### Growth and purification of P74-26 virions

P74-26 virions were prepared as previously described (18): P74-26 host strain *Thermus thermophilus HB8* (ATCC 27634) was grown overnight at 65 °C in *Thermus* growth medium (4 g L^-1^ yeast extract, 8 g L^-1^ tryptone, 3 g L^-1^ NaCl, 1 mM MgCl_2_, 0.5 mM CaCl_2_). Adsorption was initiated by combining 6 mL of fresh *T. thermophilus* culture with 4 mL of P74-26 phage stock at 1 x 10^6^ PFU/mL and incubated at 65 °C for 10 min. The adsorption reaction was inoculated into 1 L of *Thermus* medium and incubated at 65 °C while shaking for 5 hours. Phage particles were dissociated from cell debris with 1 mL chloroform and cell debris was pelleted at 4000 x g for 15 min. The supernatant was incubated with 2 U mL^-1^ DNase I and 1 μg mL^-1^ RNase A for 1 hr at 30 °C. To precipitate virions, NaCl was added to a final concentration of 1 M and PEG 8000 was added to a final concentration of 10% (w/v) while stirring, and left to incubate on ice overnight. The next morning, the phage stock was centrifuged at 11000 x g for 20 min at 4 °C. Phage pellets were dried, then resuspended in 2 mL of buffer (50 mM Tris pH 7.5, 100 mM NaCl, 1 mM MgSO_4_). 0.4 g solid CsCl was added to each resuspension and the solution was applied to a CsCl step gradient (2 mL steps each of 1.2, 1.3, 1.4, 1.5g mL^-1^ CsCl and 1 mL cushion of 1.7 g mL^-1^ CsCl, in 50 mM Tris pH 7.5, 100 mM NaCl, 1 mM MgSO_4_). The gradients were spun in a Beckman SW40-Ti rotor at 38,000 RPM for 18 hr at 4 °C. The virion layer was isolated from the gradient and dialyzed twice into 2 L of 50 mM Tris pH 8.0, 10 mM NaCl, 10 mM MgCl_2_ at 4 °C. P74-26 virions were then concentrated to 1 × 10^12^ PFU mL^-1^.

### Cloning and mutagenesis of P74-26 TTP

The P74-26 TTP (gp93) gene was *E. coli*-codon-optimized and synthesized by GenScript, then subcloned into a pSMT3 vector with a cleavable N-terminal 6x His-SUMO tag. Restriction enzymes were purchased from New England BioLabs, and oligonucleotide primers were purchased from Integrated DNA Technologies (Supplementary Table 1). Mutations were introduced using the QuikChange protocol.

### Expression and purification of P74-26 TTP

ArcticExpress (DE3) cells (Agilent) were transformed with TTP-pSMT3 and grown at 37 °C on a kanamycin (30 μg/mL) plate. A single colony was used to inoculate a 50 mL culture of 2xYT media (per liter: 16 g Bacto tryptone, 10 g yeast extract, 5 g NaCl) which was grown overnight at 37 °C in the presence of kanamycin (30 μg/mL) and gentamicin (20 μg/mL) to select for the desired plasmid and Cpn10/Cpn60 chaperonins, respectively. The overnight culture was then used to inoculate 1 L cultures which were grown at 30 °C without antibiotics for 3 hr while shaking, allowed to cool at 4 °C for 10 min, induced with a final concentration of 1 mM IPTG, then incubated at 12 °C overnight with shaking. Pelleted bacteria were then resuspended in Buffer A (50 mM Tris pH 8.0, 300 mM KCl, 20 mM imidazole, 5 mM *β*ME, 10% glycerol) and lysed by high pressure cell disruption. The lysate was centrifuged at 23000 x g for 40 min at 4 °C. The supernatant was filtered through a 0.45 μm filter (Millex, EMD Millipore) and loaded onto a 5 mL Hi-trap Ni-NTA column (Cytiva) that was pre-equilibrated in buffer A. Following lysate loading, the column was washed with 5 CV of Buffer A, then the protein was eluted in buffer B (50 mM Tris pH 8.0, 300 mM KCl, 0.5 M imidazole, 5 mM βME, 10% glycerol). 250 μL ULP1 protease was added to the eluate to cleave the tag, then dialyzed 1:1000 against Buffer A overnight. The protein was then subjected to a subtractive step over buffer A-equilibrated nickel columns to separate TTP from the tags. The protein was concentrated by centrifugal filtration (Amicon, EMD Millipore, 10-kDa MW cutoff) at 4 °C. Final protein concentration was determined by UV absorbance at 280 nm, using an extinction coefficient of 23,380 M^-1^cm^-1^.

### Electron Microscopy

#### Negative Stain EM

Carbon-coated 200 mesh copper grids (Electron Microscopy Sciences) were glow discharged on a PELCO easiGlow (Ted Pella) at 20 mA for 60 sec (negative polarity). 3.5 μL of sample was applied to the grid and incubated for 1 min. Excess sample was blotted, then the grid was washed with water followed by staining with 1% uranyl acetate (pH 4.5). Grids were viewed with a Philips CM120 electron microscope at 120 kV on a Gatan Orius SC1000 camera. Micrographs were collected between 40,000x and 66,000x.

#### Cryo-EM sample preparation

Grids were glow discharged on a PELCO easiGlow (Ted Pella) at 25 mA for 60 sec (negative polarity). *Virion tails:* 3.5 μL of purified virions at 1 × 10^10^ PFU mL^-1^ was applied to a 400-mesh C-flat holey carbon-coated grid (Electron Microscopy Sciences) at 10 °C with 90% humidity in a Vitrobot Mark IV (FEI). Sample was blotted from both sides for 10 s after a wait time of 15 s, then immediately vitrified by plunging into liquid ethane. *In vitro-assembled tubes:* 3.5 μL of purified TTP at 1.5 mg mL^-1^ was applied to a 400-mesh copper lacey carbon-coated grid (Electron Microscopy Sciences) at 10 °C with 90% humidity in a Vitrobot Mark IV (FEI). Sample was manually blotted and another 3.5 μL of sample was applied to the grid. Sample was then blotted from both sides for 8 s after a wait time of 15 s, then immediately vitrified by plunging into liquid ethane.

#### Cryo-EM data collection

Micrographs were collected on a 200 kV Talos-Arctica electron microscope (FEI) equipped with a K3 Smit direct electron detector (Gatan). *Virion tails*: Images were collected at a magnification of 45,000 in super-resolution mode with an unbinned pixel size of 0.435 Å per pixel and a total dose of 37.9644 e^-^ Å^-2^ per micrograph, with a target defocus range of −0.5 to −1.6 μm. In total, 2127 micrographs were collected. *In vitro-assembled tubes:* Images were collected as above with a total dose of 38.4312 e^-^ Å^-2^ per micrograph. In total, 2030 micrographs were collected.

#### Data Processing

Micrograph frames were aligned in IMOD with 2x binning, resulting in a pixel size of 0.87 Å per pixel. Initial CTF estimation was done using CTFFind4 within cryosparc. All following steps were done within CryoSPARC (19). *Virion tail dataset:* 217 particles were manually picked for a training dataset. From these particles, 5 pp1classes were formed and 2 were selected as templates for the filament tracer. Filament tracer was performed with a 100 *Å* filament diameter and 0.4 diameters between segments. 795,534 particles were extracted with a box size of 300 pixels and classified into 100 classes. 787,734 particles in 52 classes were used for an *ab initio* helical refinement with C1 symmetry. The resulting volume underwent consecutive rounds of symmetry searches followed by homogenous refinements until the helical parameters were determined with local search minima of a −44° twist and a 40 *Å* rise. These parameters were used along with enforced C3 symmetry in helical refinement. Local CTF refinement was performed on the 787,734 input particles and the map from the previous job was used for a final helical refinement. The Guinier plot from this job was used to obtain a B-factor value of −182.3 to sharpen the final map for a resolution of 2.73 Å. *In vitro-assembled tubes dataset:* 221 particles were manually picked for a training dataset. From these particles, 5 classes were formed and 3 were selected as templates for the filament tracer. Filament tracer was performed with a 100 Å filament diameter and 0.4 diameters between segments. 619,907 particles were extracted with a box size of 300 pixels and classified into 100 classes. 395,397 particles in 74 classes were used for a helical refinement using the helical refinement and symmetry determined above. Local CTF refinement was performed and the map was sharpened with a B-factor of −136.8 for a final resolution of 2.81 Å.

#### Model Building and Refinement

Gp93 was *de novo* built into a single subunit of the cryo-EM density of the virion tail map in Coot (33). Sidechain density was obvious due to the high resolution, so easily identifiable aromatic residues and the known sequence of gp93 were used to obtain residue register. For the model of *in vitro-* assembled tubes, the gp93 model was fitted into the *in vitro-* assembled tube map density and then manually refined. Both models were then refined into their respective maps using the Phenix real-space refine procedure (34). The Isolde plugin (35) in ChimeraX (36) was then used to further refine the map into the model followed by a final Phenix real-space refinement. The real-space refinement statistics are listed in Supplementary Table 2.

### SEC-MALS

TTP was run on a tandem size exclusion chromatography - multi-angle light scattering detector by injecting 100 μL sample at a concentration of 1 mg/mL using a 1260 Infinity HPLC (Agilent). The column was pre-equilibrated in 0.1 μm filtered buffer containing 50 mM Tris pH 8.0, 150 mM KCl, 5mM βME, 5% glycerol, and the sample was filtered through a 0.22 μm spin filter before loading. Elution was monitored by a Dawn Heleos-II MALS detector and an Optilab T-rex differential refractive index detector (Wyatt Technology). Peaks were defined and analyzed in ASTRA6 (Wyatt Technology).

### Light Scattering of tube assembly

Kinetics of tube assembly for WT and mutant TTP constructs were observed by monitoring changes in light scattering using a FluoroMax-4 spectrofluorometer (Horiba Scientific) at a wavelength of 350 nm with a 1-nm excitation bandpass and a 0.5-nm emission bandpass. Samples were loaded into a 75 μL quartz fluorometer cuvette (Starna Cells 16.50F-Q-10/Z15) with 125 μL to bring the sample meniscus above the window aperture. For each experiment, the fluorometer was programmed to equilibrate at the target temperature for 120 seconds with the cuvette in the chamber, samples were loaded from ice into the cuvette, and the program was initiated to take reads every 30 seconds with shutters closing between reads. The sample was filtered through a 0.22um filter directly prior to initiating the experiment. All experiments were performed at ~1 mg/mL unless otherwise noted.

Raw light scattering data was scaled to zero by subtracting the first value from all consecutive values so that the amplitudes of the curves could be compared directly. Light scattering curves were fit depending on the shape of the curve using GraphPad Prism version 7.04. Linear slopes were fit linearly, while non-linear slopes were fit with either a one phase association (Y=Y0 + (Plateau-Y0)*(1-exp(-K*x))) or a two phase exponential association equation (Y= amplitude1*(1-exp(-K1*X)) + amplitude2*(1-exp(-K2*X))). Curves with a sigmoidal shape were fit to (Y=Bottom + (Top-Bottom)/(1+10^((LogIC50-X)*Slope)) to account for their sigmoidal shape. The second derivative of this fit was taken in order to determine the point of inflection for sigmoidal curves, and that point was used as the location where a linear slope could be fit to the fast rate (Supplementary Figure 6).

### Electrostatic Calculations

Surface electrostatics were calculated in Pymol using the Adaptive Poisson-Boltzmann Solve (APBS) plugin (37).

### Molecular Dynamics Simulations

#### Simulation preparation

Initial structures and topology files were generated with the *tleap* module of AMBERTOOLS20 (38). All initial coordinate and topology files are included in the supplementary information. Protein interactions were described with the Amber ff19SB force field (39). Proteins were centered in a truncated octahedral periodic box solvated with the OPC four-point water model with minimum 14 Å padding (40). Sodium ions were added to make each system charge-neutral, and additional salt was added to reach 0.15 M concentration. Hydrogen mass repartitioning was applied with ParmEd (41), increasing the weight of solute hydrogen atoms to 3.024 Da. P74-26 TTP starting structures were taken from those described herein, and SPP1 TTP were taken from PDB 6YEG (12). In simulations of SPP1 TTP, C-terminal residues were found to be very flexible in absence of inter-ring interactions. Thus, for simulations of rings, we truncated residues 153-176 of the “top” ring to avoid their interaction with subunits across the periodic boundary. Further, we used additional padding (up to 22 Å) for this system. To ensure our predictions were not force field dependent, we also conducted a P74-26 TTP monomer simulation with the CHARMM36m protein force field and CHARMM-modified TIP3P water model (42). This system was initialized using the CHARMM-GUI webserver (43). For this system, the protein was centered in a rectangular periodic box with 14 Å padding, made charge-neutral and additional salt added up to 0.15 M. Hydrogen mass repartitioning was also applied to this system.

#### Simulation methodology

All simulations were performed in the gpu-accelerated *pmemd* module of the AMBER20 simulation package (44). All input files are included in the supplement. Systems were energy minimized for 500 steps of steepest descent and conjugate gradient. Systems were heated from 100 K to 310 K over 500 ps in the canonical (NVT) ensemble. Independent simulations were seeded by drawing different random initial velocities from the Maxwell-Boltzmann distribution. During heating, the temperature was controlled with the Langevin thermostat with a 1.0 ps^-1^ friction coefficient (45). A 9 Å explicit cutoff was applied, and the Particle-Mesh Ewald method was used to correct long range interactions. For the CHARMM36m monomer simulation, electrostatic interactions were calculated with a force-switching scheme, with a switching distance of 10 A and explicit cutoff of 12 A. With hydrogen mass repartitioning applied, we used a 4-fs integration timestep to propagate the equations of motion. Bonds connecting hydrogen atoms to heavy atoms were constrained with the SHAKE algorithm (46). After heating, systems were simulated in the isothermal-isobaric (NPT) ensemble at 310 K and 1 bar using the Langevin thermostat and Monte Carlo barostat (47). Simulations were run in triplicate for varying times depending on system size, with at least ~1-2 μs each for P74-26 TTP systems (Supplementary Table 5).

#### Simulation analysis

K-means clustering (*k*=10) was performed with the cpptraj program on each simulation (48). Representative frames from the largest cluster are included as PDB files in the supplement. Distances and root-meansquare fluctuations (RMSF) were calculated using cpptraj with the *distance* and *atomicfluct* commands, respectively. RMSF was calculated by first aligning the trajectories about the beta-sandwich domain of subunits (hence, loops were not included in the alignment) and then calculating a per-residue RMSF by averaging the backbone C, Cα, O, and N values. Native contacts were calculated using the *nativecontacts* command in cpptraj; the initial cryo-EM reconstructions were used as the reference structure that defined native contacts. Native contacts were calculated including only the heavy atoms in Loop1 and all the heavy atoms in the neighboring subunit(s). Supplemental movies were generated with VMD 1.9.4a55 (49).

## Supporting information

Virion Tail model coordinates

In Vitro Tube model coordinates

P74-26 monomer trajectory

P74-26 trimer trajectory

P74-26 5mer trajectory

YSD1 monomer trajectory

YSD1 11mer trajectory

SPP1 monomer trajectory

SPP1 11mer trajectory

## ACKNOWLEDGMENTS

The authors thank Drs. C. Xu, K. Song, K. Lee, and C. Ouch of the UMass Chan Medical School cryo-EM core facility for assistance with data collection, and Drs. N. Grigorieff, C. Gaubitz, N. Stone, J. Hayes, and A. Jecrois for helpful discussions and advice on data processing. We thank members of the Kelch, Royer, and Schiffer labs for helpful discussions. We thank Dr. April Pawluk of Life Science Editors and Dr. Emma Sedivy for critical reading of the manuscript. This work was funded by a National Science Foundation grant (1817338) to B. Kelch and an NSF Graduate Research Fellowship (2042597) to E. Agnello. Computational resources were provided by the NSF XSEDE Program [ACI-1053575].

## AUTHOR CONTRIBUTIONS

E.A. and B.A.K. conceptualized research. X.L. prepared the grid used for the virion tail dataset, J.P. performed molecular dynamics simulations, and E.A. performed remaining experiments. E.A., J.P., and B.A.K. analyzed data and wrote the manuscript.

The authors declare no conflict of interest.

This article is accompanied by supporting information.

## Notes

### Competing Interest Statement

The authors have declared no competing interest.

