## Supplemental Information for "Structure and assembly of an extremely long bacteriophage tail tube"

Agnello et al.

### Supporting Text

#### Unusual electrostatics of the P74-26 tail lumen

Siphovirus tails conduct DNA and encapsulated proteins from the capsid, through the lumen of the tail tube, into the infected cell. The long tail of P74-26 phage presents an interesting problem of how this molecular transfer can occur over a scale approximately an order of magnitude greater than the typical phage tail. Because the negatively charged genome must quickly traverse a long tube to allow for injection, theoretical arguments have postulated that the interior surface of tail lumens would be negatively charged such that "friction" from electrostatic interactions is minimized (1). Recent structures of sipho- and myovirus phage tails have largely upheld this view (Supplementary Figure 3) (2-6). However, we mapped the electrostatic surface of the P74-26 lumen and find it has a net neutral charge with a distribution of both positive and negative charges along the surface (Supplementary Figure 3). There also does not appear to be a presence of dyads of acidic residues as has been reported for phage tail tubes with negatively charged lumens (2-5, 7, 8). Rather, there is a relatively even distribution of acidic and basic residues. We were surprised by this finding, as we were expecting the P74-26 lumen to have similar or greater negative charge density than the lumen of phage with typical tail length. This intriguing finding suggests a range of possibilities, all of which will require more investigation. The first possibility is that a negatively charged lumen is not actually as important for ejection speed as the theoretical arguments would suggest, or that high temperature mitigates this friction. A second suggestion could be that genome ejection is slower in P74-26, which could play a functional role in its infection life cycle. A third suggestion is that the build-up of negative charges in the extraordinarily long tail decreases stability of the whole assembly, and so P74-26 has evolved a more neutral lumen to mitigate this effect. Future studies will investigate these and other possibilities for this intriguing observation.

#### Mechanisms of thermostability in self-assembling proteins

There have been numerous studies investigating thermophilic proteins, and the principles underlying their thermostability are fairly well-characterized. Although there are numerous means of stabilizing protein structure in thermophilic proteins, some general principles have emerged, such as increased reliance on hydrophobic and ionic interactions (9-14). However, structural and biochemical studies of thermophilic proteins have focused on monomeric proteins or small oligomers, therefore the principles underlying self-assembly of large polymers in thermophilic systems are still rather unexplored. Because the entropic contribution to free energy is temperature-dependent, large self-assembled systems face an extra challenge at high temperature due to their lowered entropy in the assembled state. Our study provides a step forward in elucidating the general principles underlying thermostability in self-assembling systems

A unique feature of the P74-26 tail tube is that it is constructed of trimeric rings to minimize the number of subunits. This could provide higher stability because it reduces the entropic penalty for assembly with fewer monomers needed to assemble for a given length of tube. We have observed that the P74-26 capsid likewise has a similar construction strategy in which the construction geometry minimizes the number of subunits to assemble an especially large shell (15), as does the extremophilic, archaeal virus HSTV-2 (*Halorubrum sodomense* tailed virus 2) (16). Thus, we propose that subunit minimization to enhance stability by increasing the necessity for subunit-subunit interfaces may be a widespread strategy for thermostable self-assembling systems.

Furthermore, the tail tube of phage P74-26 uses more ionic interactions to drive assembly than its mesophilic counterparts, as summarized in Supplemental Table 3 wherein we've compared the interactions in the P74-26 tail tube to those in mesophilic tail tubes SPP1, T4, and YSD1. The inter-ring interfaces, while making fewer hydrogen bonds, are substantially more electrostatic than mesophiles, making more than double the number of salt bridges on average (Supplementary Table 3). The intra-ring interfaces are substantially more hydrophobic than their mesophilic counterparts (Supplementary Table 3). Thus, it is likely that hydrophobic and ionic interactions are more stabilizing at higher temperatures, similar to what has been observed for monomeric thermophilic proteins, suggesting that this is a general strategy that all thermophilic proteins use. The preponderance of ionic interactions is also shared in the P74-26 capsid (15), and extensive hydrophobic interfaces have been shown in other hyperthermophilic viruses (17, 18).

### Supplementary Materials and Methods

#### Filament Curvature analysis

Curvature analysis in Supplemental Figure 4D was done using the kappa extension within the Fiji package of imageJ (19). A representative sample set of 15 micrographs from the virion tail and *in vitro*-assembled tube datasets each were selected. Micrograph file names were blinded to which dataset they originated, so as to avoid bias while tracing. Kappa's control point tool was used to trace all filaments in each micrograph and then calculate the average curvature of each filament.

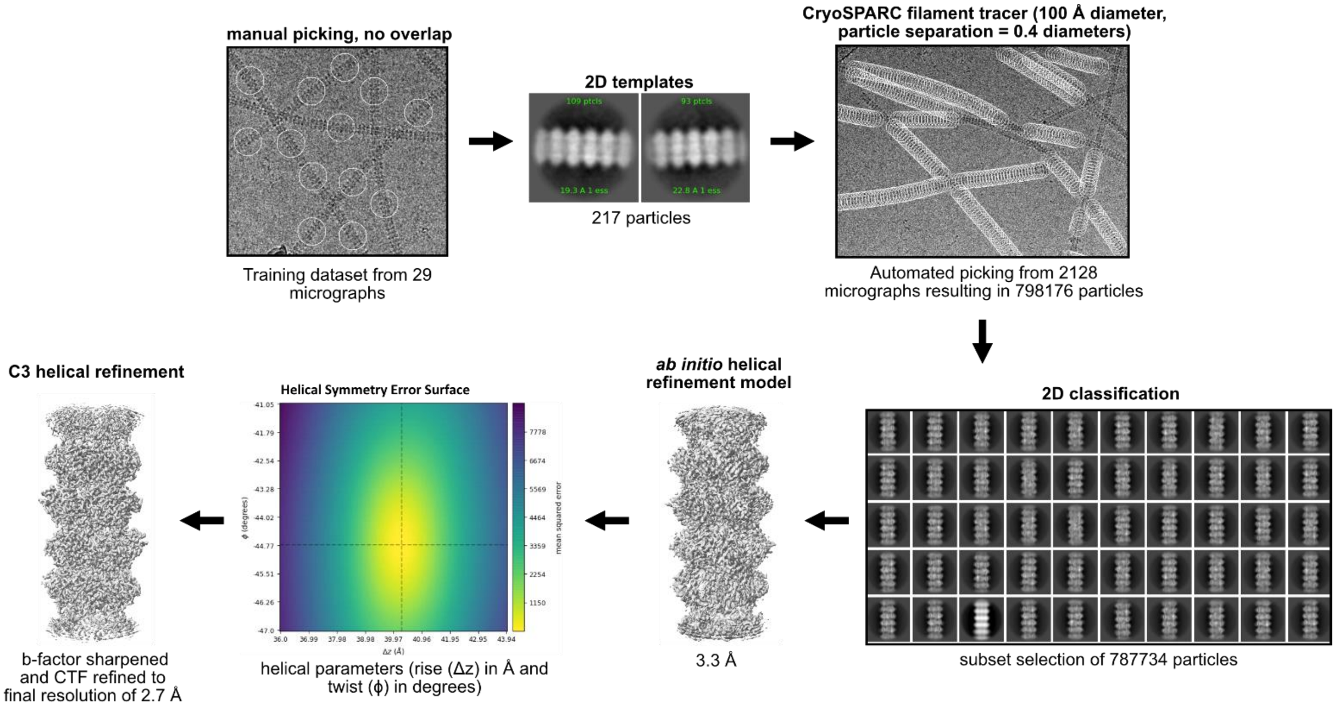

**Supplementary Figure 1. Schematic of Cryo-EM processing workflow**.
All steps shown were performed in cryoSPARC. Numbers are representative of the virion tails dataset, but the *in vitro*-assembled tubes dataset was analyzed similarly. Particles were manually picked from a subset of micrograph, then 2D classified to be used as templates for automated picking. Automated picking was done using CryoSPARC's filament tracer with a diameter of 100 Å and a particle separation of 0.4 diameters. The extracted particles were used for 2D classification of 100 classes, and 52 selected classes were used to generate an *ab initio* helical model. The helical parameters were refined using CryoSPARC's symmetry search function. The converged values were used in a helix refine job, followed by local CTF refinement and sharpening to reach a final resolution of 2.72 Å.

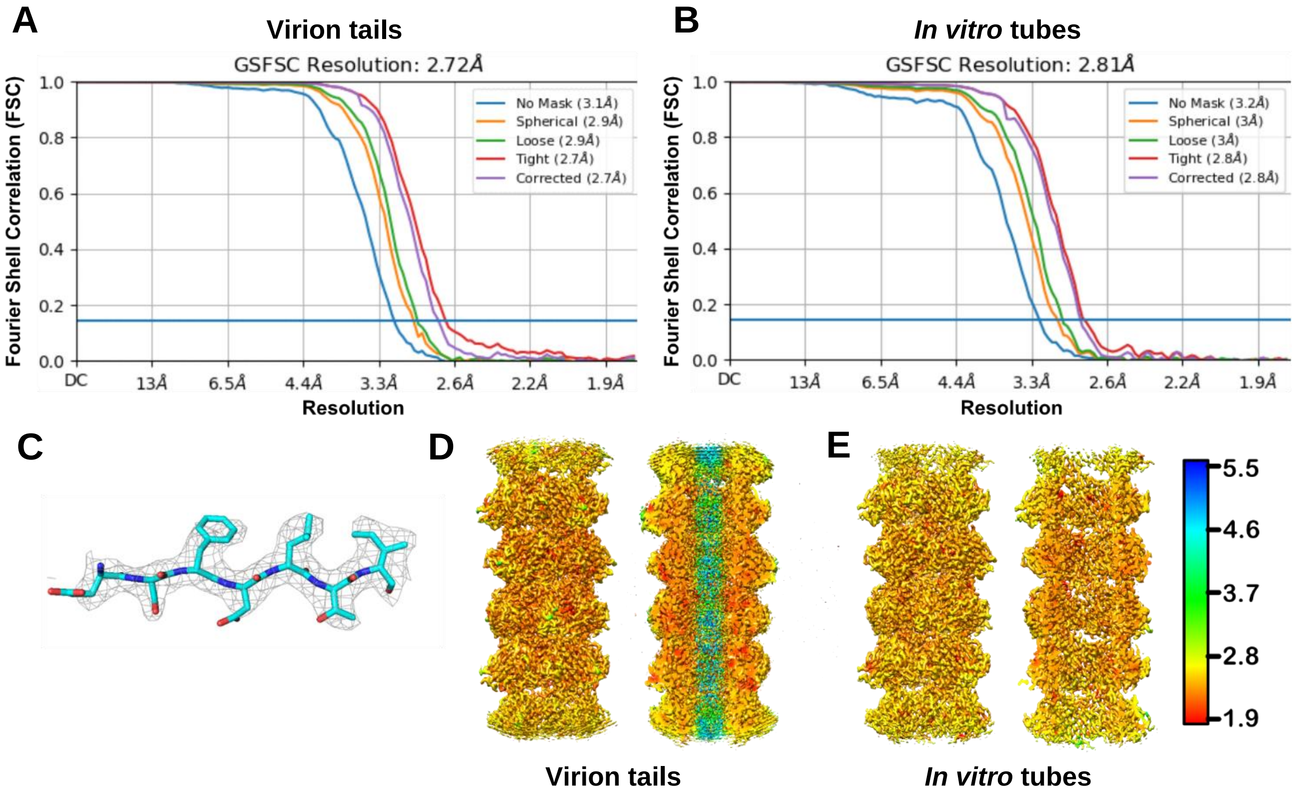

**Supplementary Figure 2. Cryo-EM structure validation and resolution**.

Gold-standard FSC curves for virion tail dataset (**A**) and *in vitro*-assembled tube dataset (**B**). Resolution was determined from the gold standard FSC cutoff of 0.143 (blue line). **C** Representative section of the virion tail density map with fitted model of TTP. **D** local resolution of maps for virion tails and **E** *in vitro*-assembled tubes from the outside of the tube (left) and an inner cutaway (right). TMP density in **D** is not well resolved.

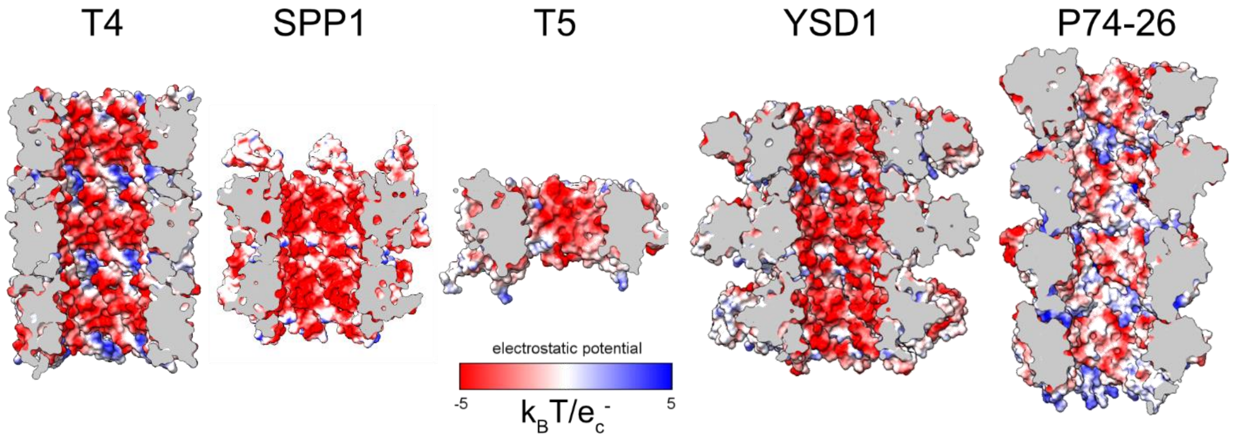

**Supplementary Figure 3. P74-26 has a uniquely neutral tail lumen.**
The interior surface of long-tailed phage is generally electronegative, thought to minimize protein-DNA interactions during genome ejection. The lumen of P74-26 is notably less electronegative than expected (see Supporting Text). PDB codes left to right: 5W5F, 6YEG, 5NGJ, 6XGR, 8ED0.

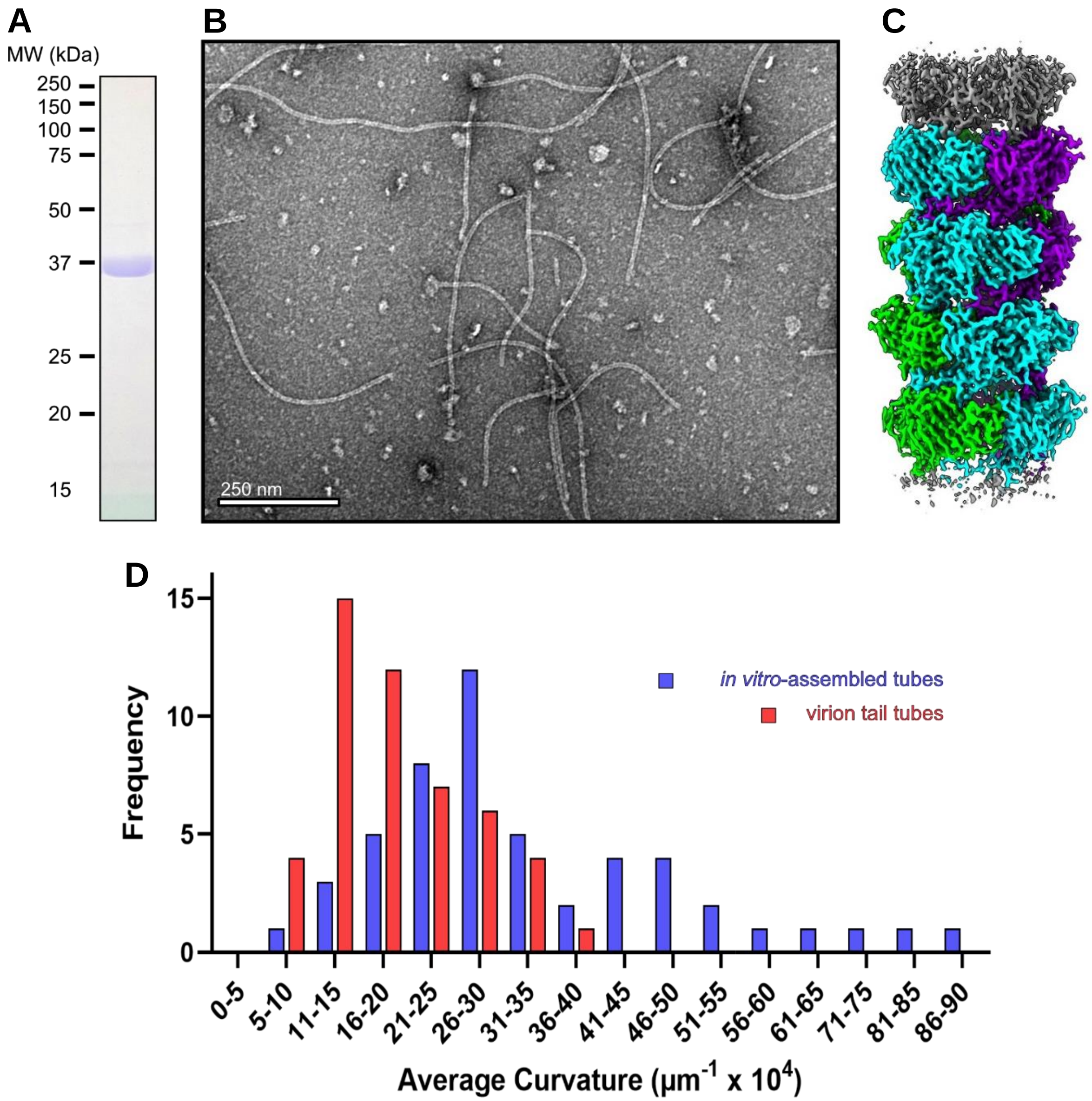

**Supplementary Figure 4. Purified tail tube protein polymerizes into flexible tail-like tubes***.*
**A** SDS-PAGE gel of TTP (37.72 kDa) after the final step of purification. **B** Negative stain EM image of purified TTP reveals spontaneous tail-like tube formation. **C** 2.8 Å reconstruction of *in vitro*-assembled TTP tubes. **D** Average filament curvature of *in vitro*-assembled TTP tubes (blue, n=51) versus virion tails (red, n=51).

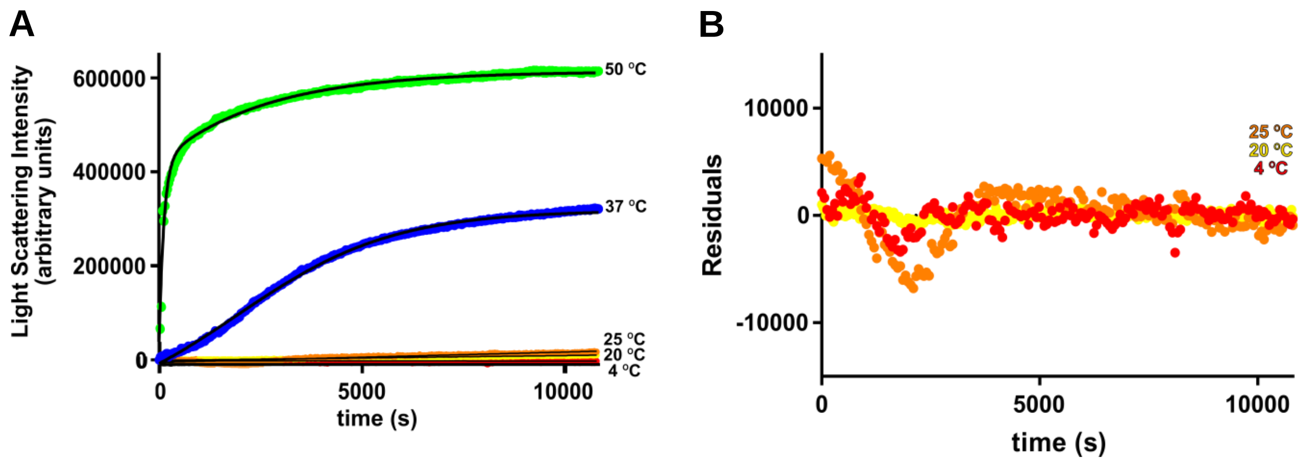

**C** **Rates of tube formation**

| rate source | Temperature (°C) | rate (LS/sec) |
|---|---|---|
| linear slope | 4 | -0.02586 |
| linear slope | 20 | 1.016 |
| linear slope | 25 | 2.195 |
| sigmoidal fast phase | 37 | 69.74 |
| k2 of two-phase | 50 | 3562 |

**Supplementary Figure 5. Raw curves and analysis of light scattering vs temperature.**
**A** Light scattering curves of TTP demonstrate the effect of temperature on TTP tube assembly. **B** Residual plot of linearly-fit data from (A). **C** Summary of rates calculated from light scattering curves. Rates were determined from the slope of linear plots (4 °C, 20 °C, 25 °C), the slope of the fast phase of sigmoidal plots (37 °C), or the majority rate of a two-phase plot (50 °C).

A

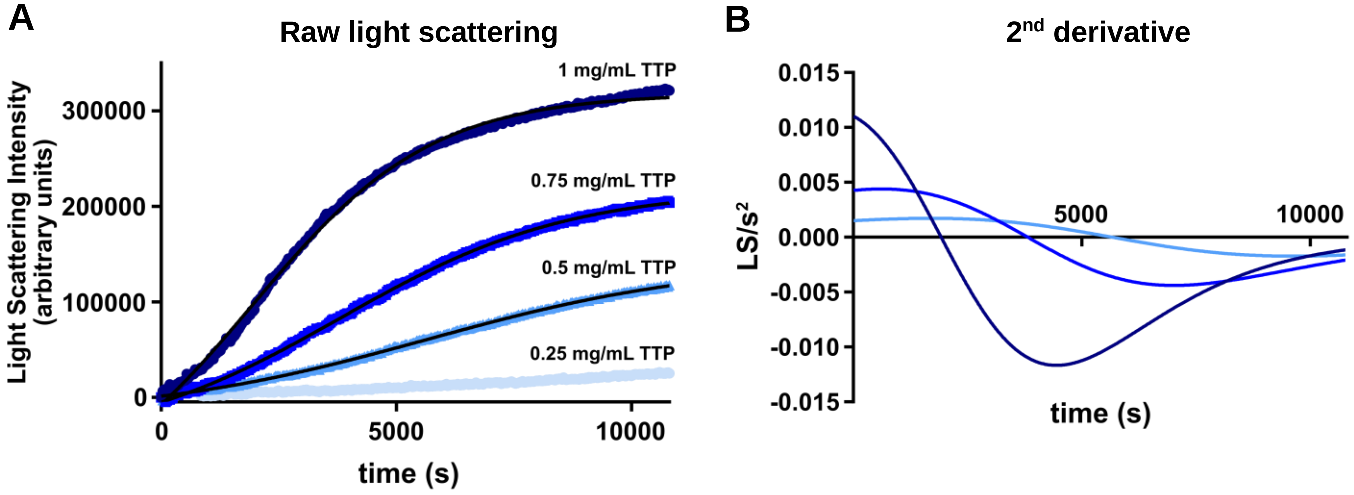

C

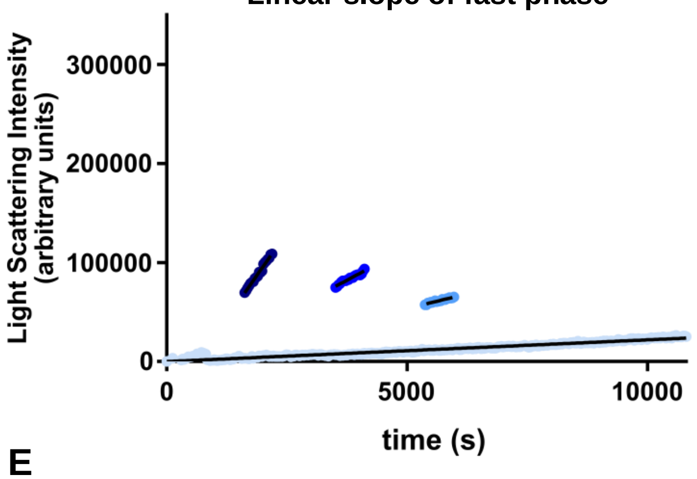

D

**Rates of tube formation**

| TTP conc (mg/mL) | Rate (LS/sec) |
|---|---|
| 1 | 69.74 |
| 0.75 | 26.07 |
| 0.5 | 11.89 |
| 0.25 | 2.322 |

E

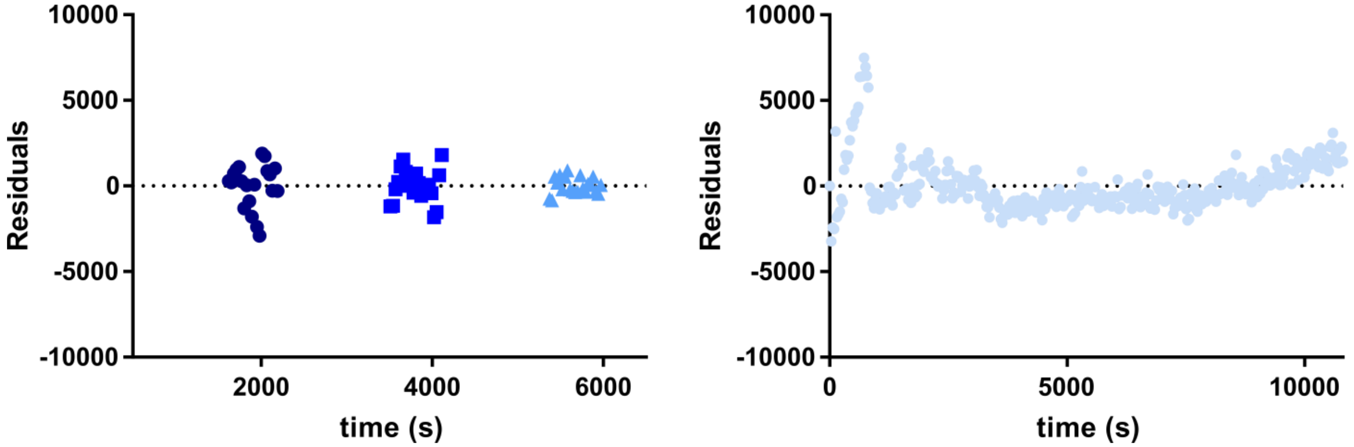

**Supplementary Figure 6. Light scattering curves and analysis for varying TTP concentrations.**
**A** Light scattering curves of WT TTP at 37 °C fitted to a sigmoidal equation (0.5-1 mg/mL). **B** The 2nd derivatives at X=0 were used to determine the inflection points of the sigmoidal curves in **(A). C** The linear slopes at the values determined in **(B)** and 0.25 mg/mL TTP. These slopes were used to determine the rates of tube formation listed in **D** in light scattering (arbitrary units) per second. **E** and **F** Residuals for the lines in **(C)** for 0.5-1 mg/mL **(E)** and 0.25 mg/mL **(F).**

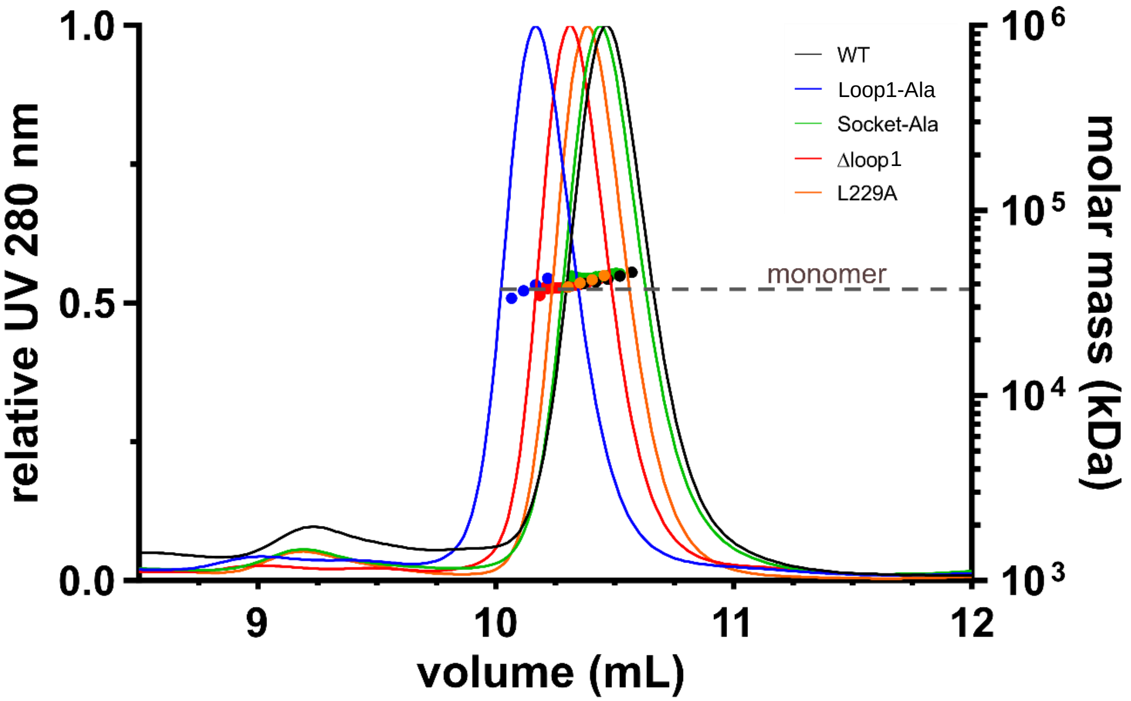

**Supplementary Figure 7. SEC-MALS of TTP Variants.**
Size Exclusion Chromatography-Multi Angle Light Scattering of TTP variants confirms their unpolymerized state is monomeric in solution.

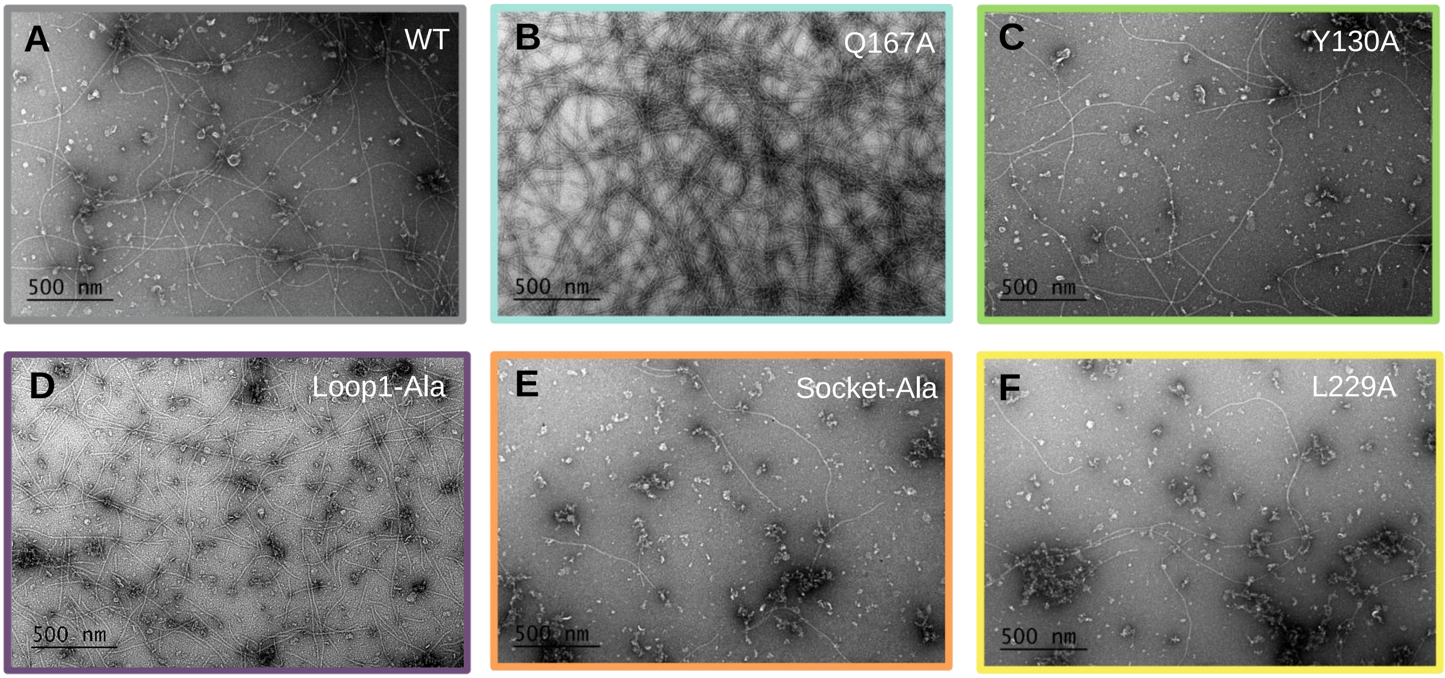

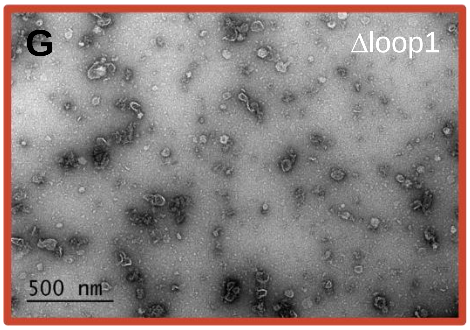

**H**

| Variant | Rate (LSU/sec) | Change relative to WT |
|---|---|---|
| WT | 1.1742 | 1x |
| Q167A | 6.7582 | 5.76x |
| Y130A | 0.3098 | 0.2638x |
| Loop1-Ala | 0.2449 | 0.2086x |
| Socket-Ala | 0.2127 | 0.1811x |
| Δloop1 | 0.1199 | 0.1021x |
| L229A | 0.07943 | 0.0676x |

**Supplementary Figure 8. Negative stain and rates of TTP variant tube formation.**
**A-G** Negative stain EM of TTP variants stained with 1% uranyl acetate, imaged at 53000x. Colored according to main text Figure 4. **H** Table of rates for tube growth, calculated from light scattering curves in main text Figure 4.

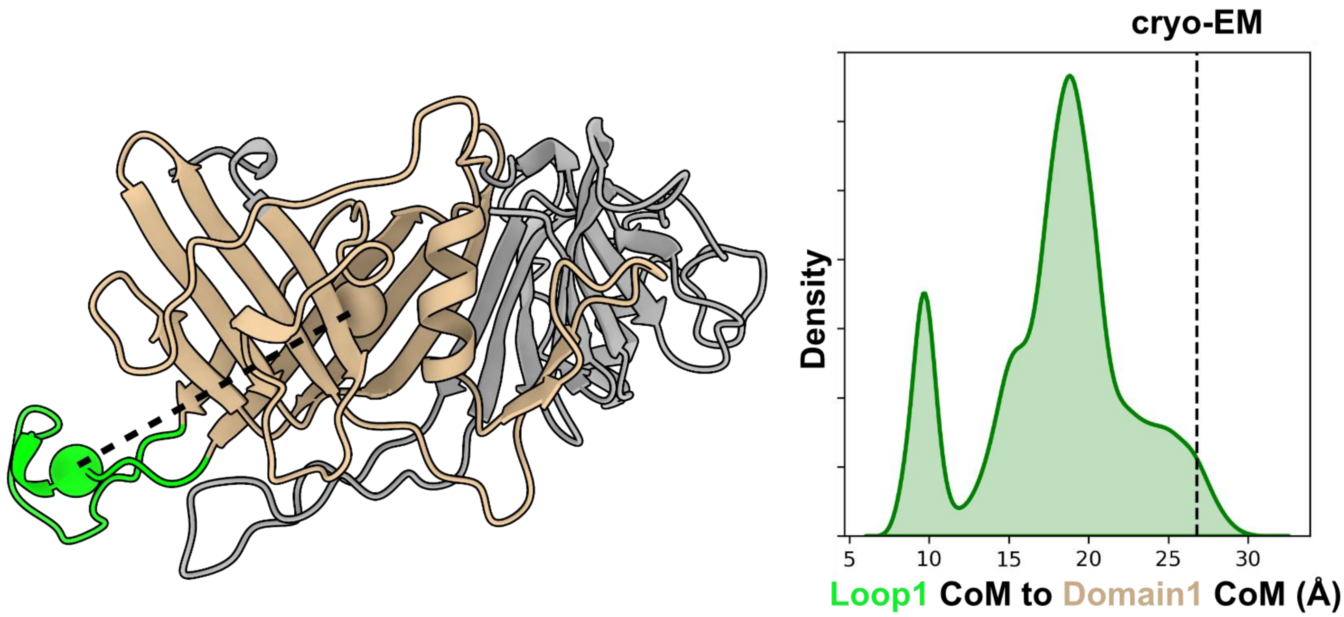

**Supplementary Figure 9. Loop1 distribution of TTP$^{P74-26}$.**

Center of mass (CoM) distribution of Loop1 of the monomer TTP$^{P74-26}$ from molecular dynamics simulations. (Left) Definitions of the CoM of Loop1 (green) and the β-sandwich domain from which it originates (tan) are shown as spheres. CoM is defined solely by the Cα of each residue. (Right) Distributions of distances sampled in molecular dynamics simulations. The dashed line represents the distance in the cryo-EM reconstruction, when assembled into a higher-order assembly.

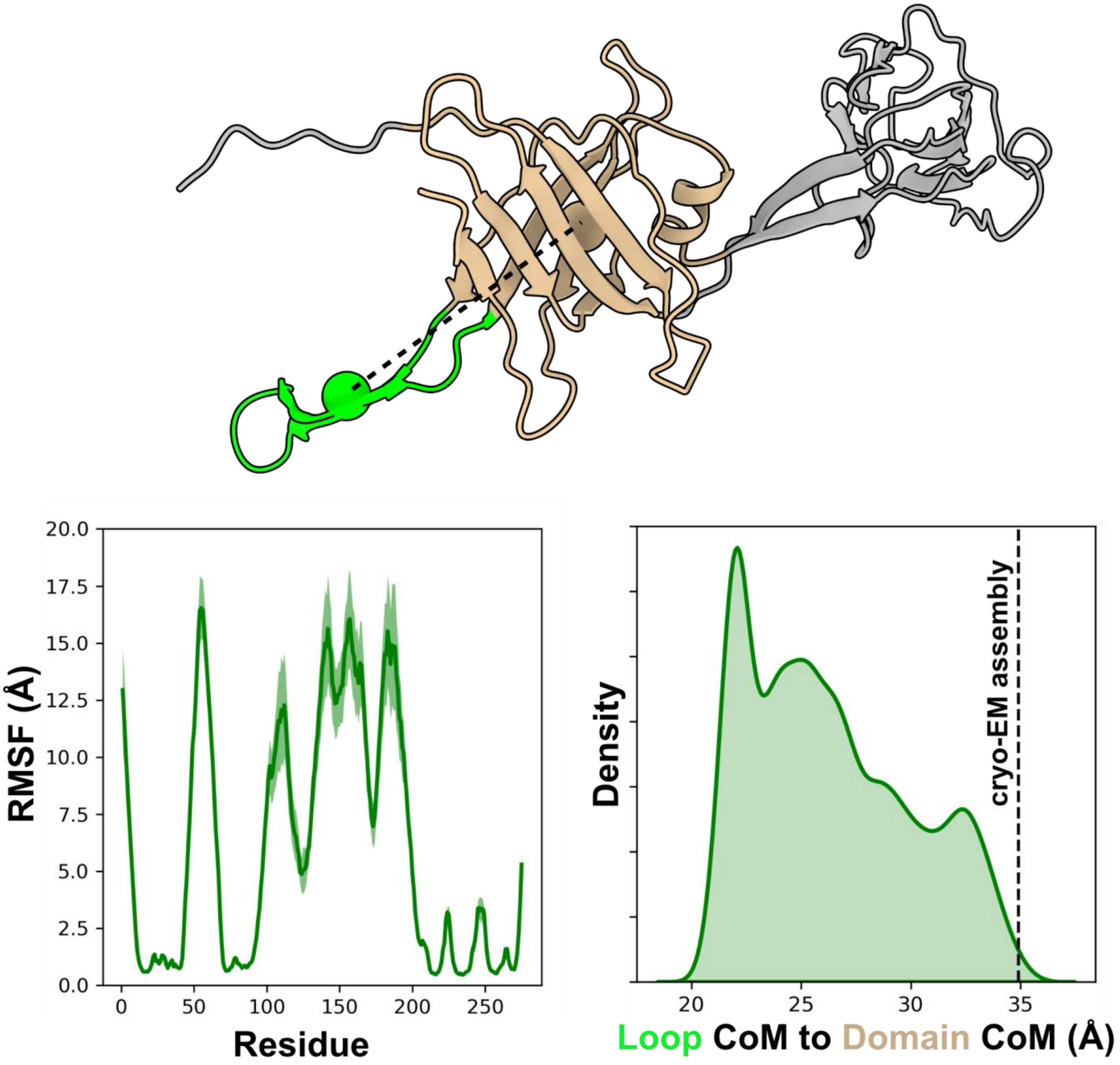

**Supplementary Figure 12. Monomer simulations of TTP$^{YSD1}$.**
(Top) Definitions of the CoM of YSD1 Loop (green) and the β-sandwich domain (tan) are shown as spheres. (Bottom left) Average Root-Mean-Square Fluctuations, showing high flexibility in the N-term, Loop, and Ig-like domain. The standard error in the mean across all replicates is shaded. (Bottom right) Distributions of distances between the Loop and β-sandwich domain sampled in molecular dynamics simulations. The dashed line represents the distance in the cryo-EM reconstruction, when assembled into a higher-order assembly.

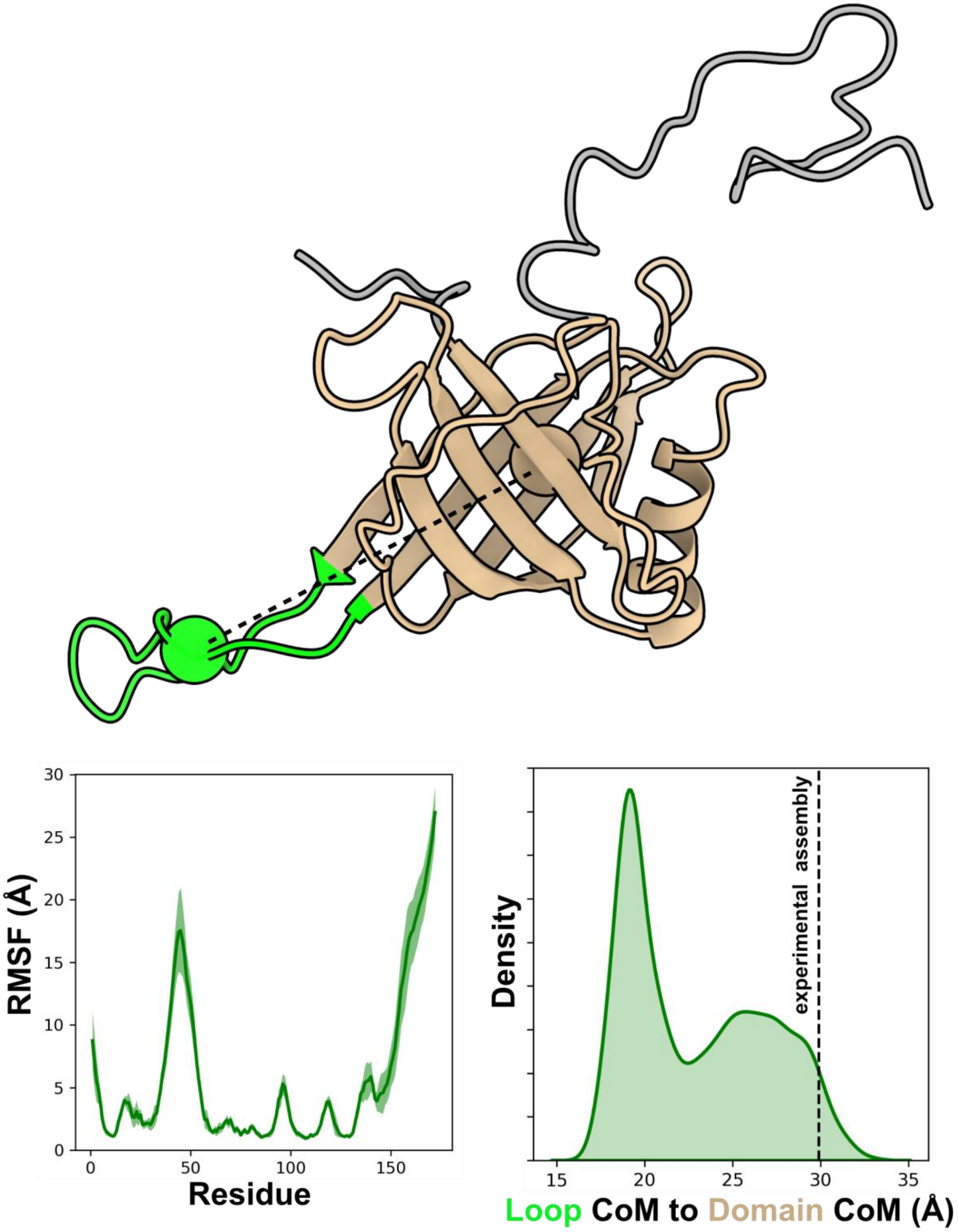

**Supplementary Figure 13. Monomer simulations of TTP$^{SPP1}$**

(Top) Definitions of the CoM of SPP1 Loop (green) and the β-sandwich domain (tan) are shown as spheres. (Bottom left) Average Root-Mean-Square Fluctuations, showing high flexibility in the N-term, Loop, and C-terminal loop. The standard error in the mean across all replicates is shaded. (Bottom right) Distributions of distances between the Loop and β-sandwich domain sampled in molecular dynamics simulations. The dashed line represents the distance in the experimental structure, when assembled into a higher-order assembly.

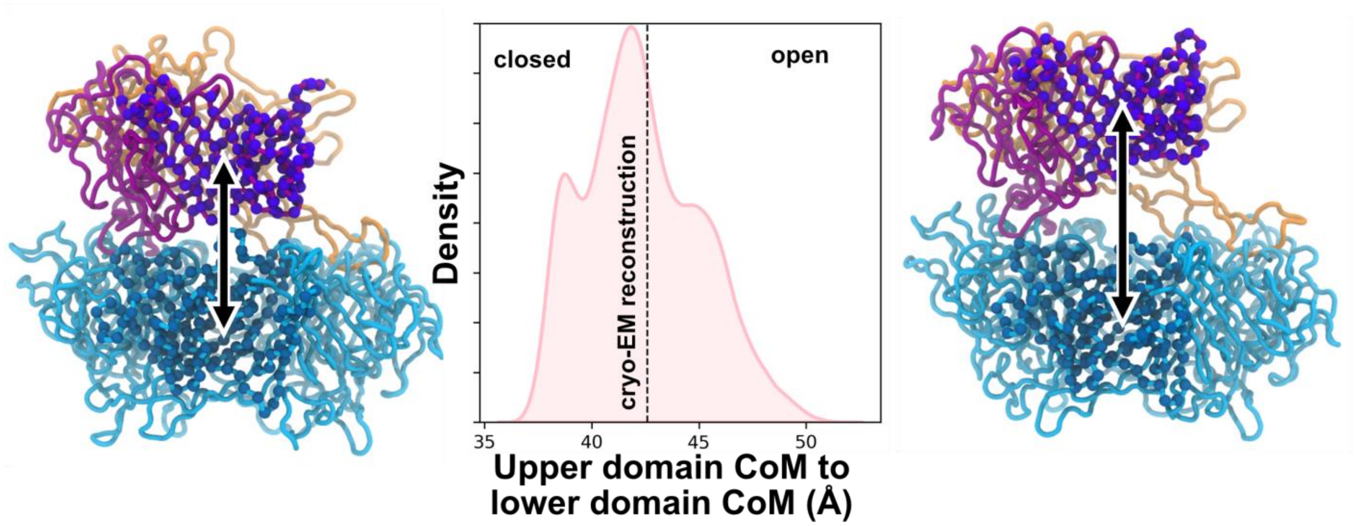

**Supplementary Figure 11. Insertion of the final subunit in a ring.**
(Center) Distance distributions between the CoM of the upper-ring subunit Domain 2 (purple) and lower ring Domain 1 (dodger blue) sampled in molecular dynamics simulations. The experimental distance of the fully assembled ring is shown as a dashed line. (Left) Example conformation of a "closed" state, where the distance between the two domains likely does not support insertion of an additional loop. (Right) Example conformation of an "open" state, where the distance between the two domains supports easy insertion of an additional loop.

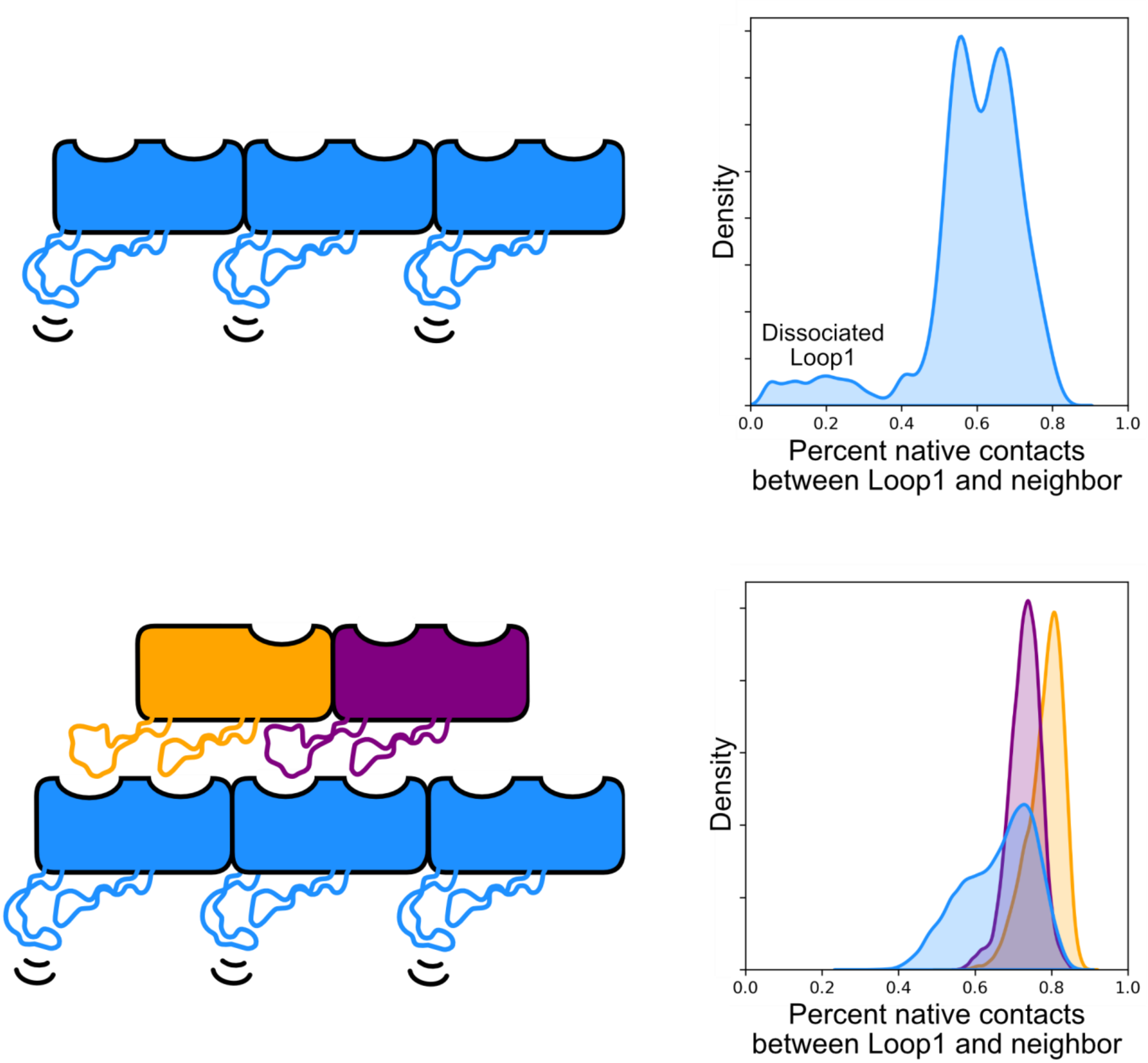

**Supplementary Figure 10. $TTP^{P74-26}$ native contact integrity sampled by molecular dynamics simulations.** (Top) The percent native contacts sampled in trimer $TTP^{P74-26}$ simulations between Loop1 of one subunit and the neighboring subunit. Native contacts are defined by the heavy atoms (non-hydrogen backbone and side chains) form the cryo-EM reconstruction, which was the starting structure for the molecular dynamics simulations. In general, intra-ring contacts alone do not completely stabilize the loop, and its flexibility leads towards loss of native contacts and potential complete dissociation of Loop1 from the neighboring subunit. (Bottom) The percent native contacts sampled in 5mer $TTP^{P74-26}$ simulations between the Loop1 of one subunit and the neighboring subunit, colored according to the subunit's unique environment. Blue subunits' Loop1 only receive intra-ring contacts, orange only receives inter-ring contacts, and purple receives both intra-ring and inter-ring contacts. In general, these inter-ring contacts (or, association of Loop1 to Socket1) are better maintained in molecular dynamics simulations. This suggests that although Loop1 remains somewhat flexible (see Main Text Figure 5), that this flexibility is accommodated by motion of Socket1 such that native contacts are preserved.

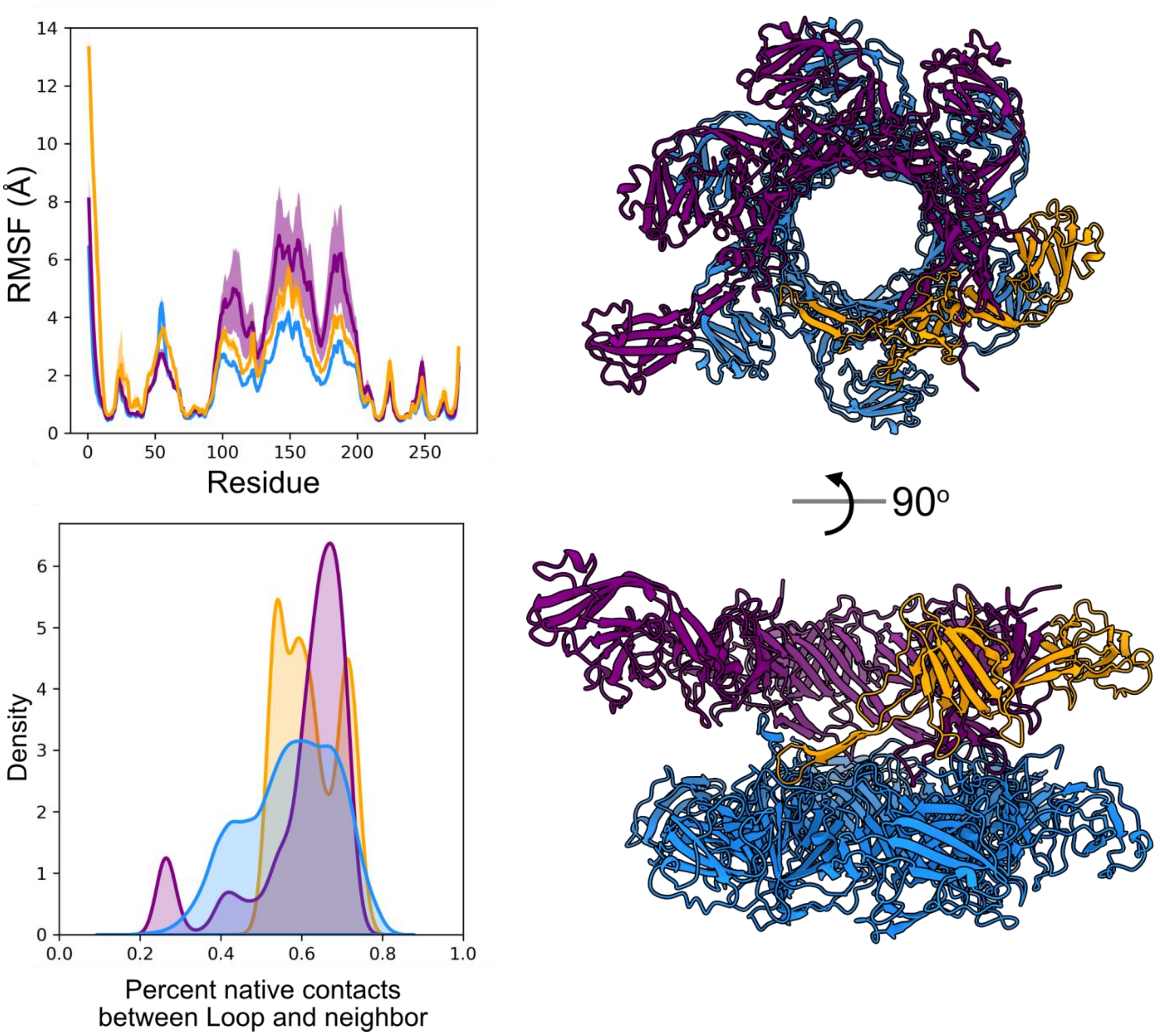

**Supplementary Figure 14. TTP$^{YSD1}$ 11mer molecular dynamics simulations.**
(Top left) Average RMSF is shown, with the standard error in the mean shaded. Like TTP$^{P74-26}$, we see that the Loops (around residue 50) are progressively stabilized by intra-ring and inter-ring contacts. (Bottom left) Percent native contacts between the Loop of one subunit and its neighboring subunits, defined similarly to that of TTP$^{P74-26}$. We note that the small peak in the purple distribution (inter-ring and intra-ring contacts) at low percentage is primarily due to a one-time buckling of the entire upper ring relative to the lower ring. (Right) A visualization of a representative structure taken from a *k=10* means clustering of a simulation.

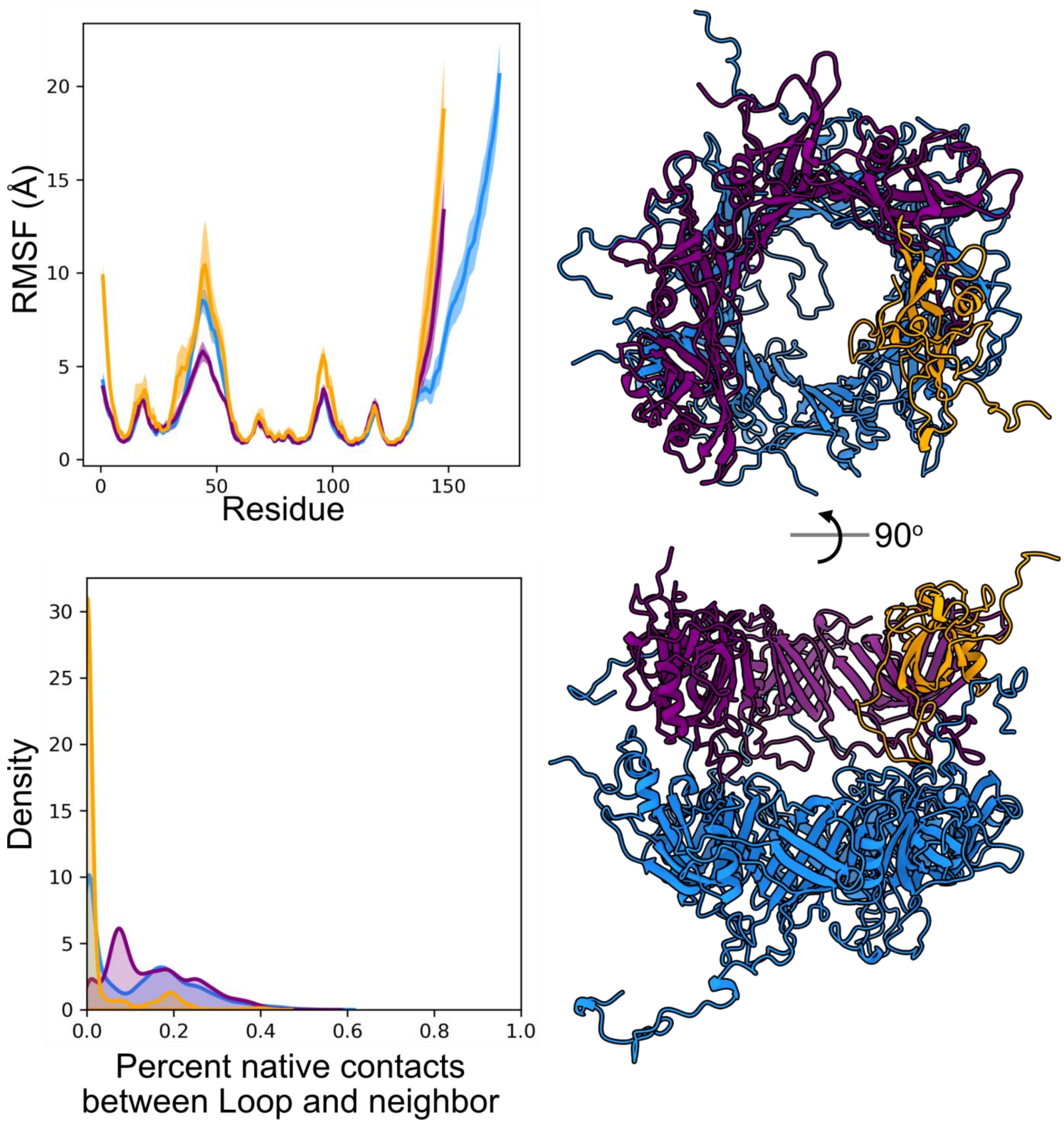

**Supplementary Figure 14. TTP$^{SPP1}$ 11mer molecular dynamics simulations.**
(Top left) Average RMSF is shown, with the standard error in the mean shaded. The Loop is primarily stabilized only by the addition of *both* inter-ring and intra-ring contacts. However, we note that the scale of this plot compared to those from either TTP$^{P74-26}$ or TTP$^{YSD1}$ suggests that this system is overall more flexible. This is reflected in the overall loss of native contacts over the course of molecular dynamics simulations (Bottom left). This could be due to several reasons; this system may require other factors to stabilize the tail tube that are not accounted for in the simulation, or the starting structure (reported resolution 4.0 Å) is unfit for molecular dynamics simulations. (Right) A visualization of a representative structure taken from a *k=10* means clustering of a simulation. It is evident that most loops have lost native contacts – including C-terminal loops from the bottom ring that extend upwards towards the top ring.

**Supplementary Movie 1: Trajectory of $TTP^{P74-26}$ monomer molecular dynamics simulation.** The trajectory was aligned about the β-sandwich domain, discounting the flexible loops from alignment.

**Supplementary Movie 2: Trajectory of $TTP^{P74-26}$ trimer molecular dynamics simulation.** The trajectory was aligned about all three subunits. We highlight some hydrophobic interactions and van der Waals spheres.

**Supplementary Movie 3: Trajectory of $TTP^{P74-26}$ 5mer molecular dynamics simulation.** The trajectory was aligned about every subunit except for the purple subunit, to highlight the opening motion over the course of the simulation.

**Supplementary Movie 4: Trajectory of $TTP^{YSD1}$ monomer molecular dynamics simulation.** The trajectory was aligned about the β-sandwich domain, discounting the flexible loops and the Ig-like domain from alignment.

**Supplementary Movie 5: Trajectory of $TTP^{SPP1}$** monomer molecular dynamics simulation. The trajectory was aligned about the β-sandwich domain, discounting the flexible loops from alignment.

**Supplementary Movie 6: Trajectory of $TTP^{YSD1}$ 11mer molecular dynamics simulation.** The trajectory was aligned about every subunit except the subunit whose Ig-like domain is not stabilized by a neighboring subunit.

**Supplementary Movie 7: Trajectory of $TTP^{SPP1}$ 11mer molecular dynamics simulation.** The trajectory was aligned about every subunit.

**Supplementary Table 1. Oligonucleotide Sequences**

| Primer Name | Primer Sequence |
|---|---|
| gp93 L229A forward | ACCGTGCGGGTAGCATTTATCGTC |
| gp93 L229A reverse | GCTACCCGCACGGTAACGGTTGGTG |
| gp93 Q167A forward | CATTCTGGCGCCGGGCGATCCGGGTTG |
| gp93 Q167A reverse | CGCCCGGCGCCAGAATGTCCTTGCC |
| gp93 Y130A forward | GGAGGGTAAGGCGGTGCGTCGTTTCTTTGGCG |
| gp93 Y130A reverse | CGACGCACCGCCTTACCCTCCGGGCCATCG |
| gp93 Socket-Ala forward | GCGGCGGCGGCGGCGCTGAACCTGAACCTGGATG |
| gp93 Socket-Ala reverse | CAGGTTCAGCGCCGCCGCCGCCGCCGCGCTGAAGGTCGCTTGGTT |
| gp93 Loop1-Ala forward | GTGATCCAGAGCGCGGCGGCGGCGGCGATGCCGATGCGTCAAATCG |
| gp93 Loop1-Ala reverse | GACGCATCGGCATCGCCGCCGCCGCCGCGCTCTGGATCACTTGCTGAC |
| gp93 Δloop1 forward | GTGGAGGCGGGTGGCGCGGTTGAAT |
| gp93 Δloop1 reverse | CTGACGACCGCTCAGGCTTTCGCTG |

**Supplementary Table 2.** Structure determination and refinement

| **Deposited Structures** | **Virion Tail** | **Tail-like tube** |
|---|---|---|
| PDB accession no. | 8ED0 | 8EDX |
| EMDB accession no. | EMD-28026 | EMD-28042 |
| **Data Collection** | | |
| Microscope | FEI Talos Arctica | FEI Talos Arctica |
| Detector | Gatan K2 | Gatan K2 |
| Voltage (kV) | 200 | 200 |
| Magnification | 45000 | 45000 |
| Electron exposure ($e^-$/Å$^2$) | 37.9644 | 38.4312 |
| Defocus range (µm) | -0.5 to -1.5 | -0.5 to -1.5 |
| Pixel size (Å) | 0.435 | 0.435 |
| **Data Processing** | | |
| Number of particles refined | 795,534 | 619,907 |
| Final number of particles | 787,734 | 395,357 |
| Imposed Symmetry | C3 | C3 |
| Map-sharpening B-factor (Å$^2$) | -150.5 | -136.8 |
| Final Resolution (Å, FSC 0.143) | 2.72 | 2.81 |
| **Subunit Refinement** | | |
| Map Correlation (%) (CC_mask) | 0.90 | .91 |
| R.M.S.D. (bonds) | 0.006 | .006 |
| R.M.S.D. (angles) | 0.720 | .557 |
| All-atom Clash score | 5.07 | 2.63 |
| Ramachandran favored (%) | 95.66 | 95.38 |
| Ramachandran allowed (%) | 3.76 | 4.63 |
| Ramachandran outliers (%) | 0.58 | 0.00 |
| Rotamer outliers (%) | 0.00 | 0.35 |
| C-beta deviations | 0.00 | 0.00 |

**Supplementary Table 3. PISA Interface Analysis of Bacteriophage Tail Tubes**

| Phage name (PDB) | Interface Area (Å$^2$) | ΔG (kcal/mol) | ΔG P-value | $N_{HB}$ | $N_{SB}$ | $N_{DS}$ | Fraction of total interface area | Scaled ΔG p-value* |
|---|---|---|---|---|---|---|---|---|
| **SPP1** (6YEG) | | | | | | | | |
| Intra-ring (average) | 1372 | -6.50 | 0.724 | 17 | 1 | 0 | 1.000 | 0.724 |
| | | | | | | | | total: 0.724 |
| Inter-ring | 856.2 | -12.8 | 0.189 | 3 | 0 | 0 | 0.539 | 0.102 |
| | 599.8 | -2.20 | 0.708 | 4 | 2 | 0 | 0.377 | 0.267 |
| | 98.90 | -1.70 | 0.292 | 0 | 0 | 0 | 0.0622 | 0.0182 |
| | 34.30 | 1.20 | 0.839 | 1 | 0 | 0 | 0.0216 | 0.0181 |
| | | total per hexameric ring: | | 48 | 12 | 0 | | total: 0.405 |
| **T4** (5W5F) | | | | | | | | |
| Intra-ring (average) | 2149 | -16.4 | 0.560 | 27 | 1 | 0 | 0.947 | 0.530 |
| | 119.7 | -0.600 | 0.588 | 3 | 0 | 0 | 0.0528 | 0.0310 |
| | | | | | | | | total: 0.561 |
| Inter-ring | 392.3 | -5.00 | 0.213 | 3 | 0 | 0 | 0.393 | 0.0837 |
| | 296.3 | -2.20 | 0.523 | 0 | 0 | 0 | 0.297 | 0.155 |
| | 169.4 | -2.00 | 0.381 | 4 | 0 | 0 | 0.170 | 0.0647 |
| | 140.3 | -1.50 | 0.448 | 0 | 0 | 0 | 0.141 | 0.0630 |
| | | total per hexameric ring: | | 42 | 0 | 0 | | total: 0.367 |
| **YSD1** (6XGR) | | | | | | | | |
| Intra-ring (average) | 2519 | -18.7 | 0.456 | 33 | 11 | 0 | 0.953 | 0.434 |
| | 124.8 | 0.400 | 0.746 | 1 | 0 | 0 | 0.0472 | 0.0352 |
| | | | | | | | | total: 0.470 |
| Inter-ring | 566.4 | -4.20 | 0.429 | 4 | 2 | 0 | 0.714 | 0.306 |
| | 165.4 | -2.30 | 0.337 | 2 | 0 | 0 | 0.208 | 0.0702 |
| | 61.90 | -0.400 | 0.731 | 1 | 0 | 0 | 0.0780 | 0.0570 |
| | | total per hexameric ring: | | 42 | 12 | 0 | | total: 0.433 |
| **P74-26** | | | | | | | | |
| Intra-ring (average) | 1487 | -14.3 | 0.179 | 7 | 4 | 0 | 1.000 | 0.179 |
| | | | | | | | | total: 0.179 |
| Inter-ring | 1124.4 | -8.20 | 0.371 | 5 | 10 | 0 | 0.891 | 0.331 |
| | 77.30 | -2.20 | 0.114 | 0 | 0 | 0 | 0.0613 | 0.00699 |
| | 59.70 | 1.10 | 0.784 | 0 | 0 | 0 | 0.0473 | 0.0371 |
| | | total per trimeric ring: | | 15 | 30 | 0 | | total: 0.375 |

*Protein interfaces, surfaces and assemblies' service PISA at the European Bioinformatics Institute* (20)

Inter-ring interactions of P74-26 have less hydrogen-bonds (red), but more electrostatics (green), compared to mesophilic phage tails. Intra-ring interactions of P74-26 are more hydrophobic than mesophiles, indicated by a lower ∆G P-value (blue). ∆G P-value indicates the P-value of the observed solvation free energy gain, or the probability of getting a lower than observed $\mathbf{\Delta^{i}G}$, when the interface atoms are picked randomly from the protein surface. **P<0.5** indicates interfaces with higher than would-be-average hydrophobicity, implying that the interface surface can be interaction-specific.

*Scaled P-values are calculated by multiplying the raw ∆G P-value by the fraction of total interface area.

**Supplementary Table 4.**
**PISA Interface Analysis of P74-26 Tail Tube**

| Interface | Interface Area (Å$^2$) | ΔG (kcal/mol) | $N_{HB}$ | $N_{SB}$ | $N_{DS}$ |
|---|---|---|---|---|---|
| Intra-ring | 1486.5 | -14.3 | 7 | 4 | 0 |
| Inter-ring | 1124.4 | -8.2 | 5 | 10 | 0 |
| | 77.3 | -2.2 | 0 | 0 | 0 |
| | 59.7 | -1.1 | 0 | 0 | 0 |
| | total: 1261.4 | | | | |

**PISA Interface Analysis of P74-26 Tail Tube with Loop1 only**

| Interface | Interface Area (Å$^2$) | ΔG (kcal/mol) | $N_{HB}$ | $N_{SB}$ | $N_{DS}$ |
|---|---|---|---|---|---|
| Intra-ring | 1429.6 | 13.6 | 6 | 4 | 0 |
| Inter-ring | 720.1 | -3.8 | 3 | 6 | 0 |
| | 77.3 | -2.2 | 0 | 0 | 0 |
| | 14.5 | 0.1 | 0 | 0 | 0 |
| | total: 811.9 | | | | |

**PISA Interface Analysis of P74-26 Tail Tube with Loop2 only**

| Interface | Interface Area (Å$^2$) | ΔG (kcal/mol) | $N_{HB}$ | $N_{SB}$ | $N_{DS}$ |
|---|---|---|---|---|---|
| Intra-ring | 839.0 | -7.1 | 6 | 4 | 0 |
| Inter-ring | 461.9 | -5.3 | 2 | 4 | 0 |
| | 59.7 | 1.1 | 0 | 0 | 0 |
| | total: 521.6 | | | | |

**PISA Interface Analysis of P74-26 Tail Tube with both loops deleted (42-65, 220-242)**

| Interface | Interface Area (Å$^2$) | ΔG (kcal/mol) | $N_{HB}$ | $N_{SB}$ | $N_{DS}$ |
|---|---|---|---|---|---|
| Intra-ring | 840.9 | -7.2 | 6 | 4 | 0 |
| Inter-ring | 14.5 | 0.1 | 0 | 0 | 0 |
| | 10.4 | 0.2 | 0 | 0 | 0 |
| | total: 24.9 | | | | |

**Supplementary Table 5.** Summary of simulations

| System | Force Field | Simulation Time (μs) |
|---|---|---|
| P74-26 monomer (replicate 1) | ff19SB | 1.99 |
| P74-26 monomer (replicate 2) | ff19SB | 2.01 |
| P74-26 monomer (replicate 3) | ff19SB | 2.01 |
| P74-26 monomer (replicate 4) | C36m | 2.01 |
| P74-26 trimer (replicate 1) | ff19SB | 1.6 |
| P74-26 trimer (replicate 2) | ff19SB | 1.69 |
| P74-26 trimer (replicate 3) | ff19SB | 1.7 |
| P74-26 5mer (replicate 1) | ff19SB | 1.15 |
| P74-26 5mer (replicate 2) | ff19SB | 1.15 |
| P74-26 5mer (replicate 3) | ff19SB | 1.15 |
| SPP1 monomer (replicate 1) | ff19SB | 0.96 |
| SPP1 monomer (replicate 2) | ff19SB | 0.97 |
| SPP1 monomer (replicate 3) | ff19SB | 0.56 |
| SPP1 11mer (replicate 1) | ff19SB | 0.43 |
| SPP1 11mer (replicate 2) | ff19SB | 0.43 |
| SPP1 11mer (replicate 3) | ff19SB | 0.43 |
| YSD1 monomer (replicate 1) | ff19SB | 1.24 |
| YSD1 monomer (replicate 2) | ff19SB | 1.25 |
| YSD1 monomer (replicate 3) | ff19SB | 1.24 |
| YSD1 11mer (replicate 1) | ff19SB | 0.75 |
| YSD1 11mer (replicate 2) | ff19SB | 0.6 |
| YSD1 11mer (replicate 3) | ff19SB | 0.59 |
